# Novel Ser74 of NF-κB/*Cg*IκBα Phosphorylated by MAPK/ERK Regulates Temperature Adaptation in Oysters

**DOI:** 10.1101/2024.03.11.584362

**Authors:** Chaogang Wang, Zhuxiang Jiang, Mingyang Du, Rihao Cong, Wei Wang, Taiping Zhang, Jincheng Chen, Guofan Zhang, Li Li

## Abstract

Phosphorylation of IκBα at Ser32 and Ser36 by IKKs during biotic stress triggers its ubiquitin-proteasome degradation, causing to the nuclear translocation of REL, representing a key cascade mechanism in metazoans conserved and immune core signaling pathway, NF-κB. However, studies on its response to abiotic stress and signal transduction by phosphorylation in mollusks are lacking. Here, we firstly report a novel heat-induced phosphorylation site (Ser74) at the major NF-κB/*Cg*IκBα of oysters, phosphorylated by MAPK/*Cg*ERK1/2, which independently mediated the subsequent ubiquitin-proteasome degradation without phosphorylation at Ser32 and Ser36 and decreased thermal stability. The degradation of *Cg*IκBα promoted *Cg*REL nuclear translocation, which stimulated cell survival related gene expression to defend against thermal stress. The MAPK and NF-κB pathways exhibited stronger activation patterns in higher environmental temperature and in the warm-adapted *Crassostrea angulata* than those in the cold-adapted *C. gigas*-two allopatric congeneric oyster species with differential habitat temperatures. These findings unveil the complex and unique phosphorylation-mediated signal transduction mechanisms in marine invertebrates, and further expand our understanding of the evolution and function of established classical pathway crosstalk mechanisms.

## Introduction

Studying the regulatory mechanism underlying temperature adaptation is crucial to understanding the impact and adaptive potential of the organism in response to climate change. Protein phosphorylation is a widely studied post-translational modification that regulates protein localization, influences protein–protein interactions, and participates in protein degradation, making it crucial in mediating stress signal transduction under temperature stress in vertebrates (Yoshihara, Naito et al., 2013), plants (Kamal, Ishikawa et al., 2020, Pang, Hu et al., 2021), *Caenorhabditis elegans* (Huang, Wu et al., 2020), *Drosophila melanogaster* (Elvira, Cha et al., 2020), and *Saccharomyces cerevisiae* (Kanshin, Kubiniok et al., 2015). Temperature is a fundamental abiotic stress factor, which can directly influence metabolic rates, enzyme activity, membrane fluidity, and cellular functions, which activates numerous conserved signaling pathways by mobilizing corresponding kinases and triggering signal cascades through phosphorylation, including AMPK-mTOR (Gonzalez, Hall et al., 2020, Zhao, Hu et al., 2017), PI3K-AKT (Oehler-Jänne, Bueren et al., 2008), NF-κB (Harper, Woodcock et al., 2018, Paszek, Kardyńska et al., 2020), and MAPK (Moustafa, AbuQamar et al., 2014) , etc. Studies on phosphorylation in response to temperature stress in mollusks are lacking, as they are currently limited to the measurement of phosphorylation levels of specific proteins within specific pathways, such as SAPK/JNK (Evans & Somero, 2010) or p38 MAPK (Michaelidis, Hatzikamari et al., 2009) in MAPK pathway, p53 and p21 (Yao & Somero, 2013). To our knowledge, only our previous study has investigated the global phosphoproteomic response to high-temperature stress in mollusks (oysters), revealing a significant upregulation of phosphorylation level at the Ser74 site of an NF-κB/IκB protein under high-temperature stress, which exhibited divergent phosphorylation patterns between two differentially heat-adaptive species (Wang, Du et al., 2023a), while the conserved Ser32 and Ser36 sites showed no phosphorylation modification, suggesting a potential novel temperature-induced phosphorylation regulatory mechanisms in the NF-κB pathway for oysters.

The NF-κB signaling pathway is essential in organisms and crucial in regulating numerous physiological processes, including development, inflammation, cell apoptosis, cell proliferation, differentiation, and immune response (Ghosh & Hayden, 2008, Hayden & Ghosh, 2008, Sun & Ley, 2008). In mammals, the NF-κB/Rel family members can be divided into two subfamilies based on their C-terminal structures: the NF-κB and Rel subfamilies. The Rel subfamily comprises three members: c-Rel, RelA (p65), and RelB, which have their C-terminal domains responsible for transcriptional activation (Hayden & Ghosh, 2012, Siebenlist, Franzoso et al., 1994, Zheng, Yin et al., 2011). The inhibitors NF-κB (IκB) are considered crucial regulatory factors of the NF-κB signaling pathway (Hinz, Arslan et al., 2012). Its core signaling cascade involves the stimulation of IKK complex (comprising IKKβ, IKKα, and NEMO) kinases by pro-inflammatory cytokines, LPS, growth factors, and antigen receptors, phosphorylating IκB (a key regulatory factor in the NF-κB signaling pathway) at the Ser32 and Ser36 sites, which is conserved across various metazoans and can promote its ubiquitination-proteasome degradation (Chen & Chen, 2013, Hayden & Ghosh, 2008, Hinz et al., 2012, Perkins, 2006). These events cause the nuclear translocation of REL and trigger the activation of downstream gene expression (Chen & Chen, 2013, Perkins, 2006). Over the past decade, the role of NF-κB, which is essential in the antiviral and antibacterial processes for vertebrate and invertebrate cells, has been focused on the study of animal immune defense (Dong, Sang et al., 2020, Ghosh & Hayden, 2008, Goodson Michael, Kojadinovic et al., 2005, Hayden & Ghosh, 2008, Hinz et al., 2012, Kasthuri, Whang et al., 2013, Khush, Leulier et al., 2001, Li, Wang et al., 2019, Oyanedel, Gonzalez et al., 2016, Sang, Dong et al., 2020, Xu, Li et al., 2015, Zhang, Jiang et al., 2009). And current researches has mainly concentrated on its roles in immune diseases (Li & Verma, 2002, Liu, Zhang et al., 2017), inflammation regulation (Liu et al., 2017), and cancer progression (Karin, 2006) in mammals and arthropods (such as *D. melanogaster*) (Khush et al., 2001, Sun & Ley, 2008), lacking reports about its response to abiotic stressors, such as temperature. However, several studies have proven that the NF-κB signaling pathway can also be activated by stimuli such as free radicals, reactive oxygen species, and cytokines, which are typical products of temperature stress (Moniruzzaman, Ghosal et al., 2018). Additionally, current studies on phosphorylation-mediated NF-κB regulatory mechanisms in mollusks focus primarily on identifying Rel/NF-κB and IκB homologs and their expression response during biotic stress in Pacific oyster (*Crassostrea gigas*) (Dong et al., 2020, Montagnani, Labreuche et al., 2008, Sang et al., 2020, Xu et al., 2015), scallop (*Argopecten purpuratus*) (Oyanedel et al., 2016), sea sleeve (*Euprymna scolopes*) (Goodson Michael et al., 2005), abalone (*Haliotis discus discus*) (Kasthuri et al., 2013) and pearl oyster (*Pinctada fucata*) (Zhang et al., 2009). Therefore, elucidating the function of phosphorylation of novel Ser74 site at IκB, upstream kinase and signaling cascade pathway mediating its phosphorylation, and downstream REL-activated genes during high temperature will bridge the gap of the phosphorylation-mediated signal transduction for the marine invertebrate in response to the stress and will facilitate understanding the importance and complexity of phosphorylation regulation in temperature adaptation.

Oysters, as representative species with early deciphered genomes in mollusks, are distributed worldwide and possess significant economic and ecological values (Guo, 2009, Zhang, Fang et al., 2012). Furthermore, oysters inhabit the excessively stressful intertidal zone, which has caused the evolution of perfect tolerance to such conditions (Clark, Thorne et al., 2013), making them an ideal model organism for revealing the regulatory mechanisms underlying stress adaptation mediated by specific signaling pathways in mollusks. *Crassostrea gigas* (*C. gigas*) and *Crassostrea angulata* (*C. angulata*) are two allopatric congeneric oyster species that adapt to relatively cold and warm habitats (Northern and Southern China coasts), respectively, exhibiting divergent temperature adaptation (Ghaffari, Wang et al., 2019, Haiyan, Guofan et al., 2008, Ren, Liu et al., 2010, Wang, Li et al., 2023b, Wang, Li et al., 2021). Previous study has found differential protein phosphorylation patterns in response to high-temperature stress between *C. gigas* and *C. angulata* (Wang et al., 2023a). Particularly, opposite phosphorylation levels have been observed in IκB (Ser74) suggesting that NF-κB/IκBα phosphorylation may be crucial in shaping divergent temperature adaptation between the oyster species. Therefore, comparative studies between these two species will facilitate revealing the phosphorylation mechanism of the NF-κB/IκB (Ser74) pathway and its cascade network in mollusks and the role of its phosphorylation regulation in shaping divergent temperature adaptation.

In this study, we discovered a novel heat-induced phosphorylation site (Ser74) at major *Cg*IκBα of immune core NF-κB pathway, which had a distinct function independent of the Ser32 and Ser36 sites in mediating its ubiquitin-proteasome degradation and thermal stability, through phylogenetic analysis, sequence alignment, and molecular experiments. Further functional validation experiments confirmed that this site was phosphorylated by MAPK/*Cg*ERK1/2 kinase, and activated cell survival, fatty acid metabolism, protein translation, and antioxidant gene expression by promoting *Cg*REL nuclear translocation, thereby helping organisms withstand heat stress. And this pathway was regulated by the classical MAPK pathway through the phosphorylation. From MAPK to NF-κB pathways, the phosphorylation and expression levels of key kinases, regulatory factors, and downstream activated genes all exhibit differential temperature responses and adaptation patterns, which suggests their involvement in shaping the divergent temperature adaptation between *C. gigas* and *C. angulata*, two allopatric congeneric oyster species with differential habitat temperatures. Our findings reveal the existence of complex and unique phosphorylation-mediated signal transduction mechanisms in marine invertebrates, such as oysters, and expand the understanding of the evolution and function of the established classical pathways crosstalk in temperature adaptation.

## Results

### Specific evolution of Ser74 in oyster’s major IκBα

The phylogenetic analysis of the IκB protein family across vertebrates and invertebrates revealed that marine mollusks possess only BCL3 and IκBα proteins, accompanied by variations in the number of ANK domains (Table S1, Fig. 1A). Notably, the IκBα protein demonstrated an expansion in copy number in Bivalvia and Gastropoda, such as *C. gigas* had four IκBα proteins. Expression analysis of *C. gigas* demonstrated distinct variations among the four IκBα genes, with *Cg* IκB like4 exhibiting significantly higher expression than the other three IκBα genes. Moreover, the expression of *Cg* IκB like4 displayed tissue specificity, with marked higher expression observed in the gill tissue and hemocyte (Fig. 1B). Insignificant changes were noted in the expression across different developmental stages.

**Fig. 1.**
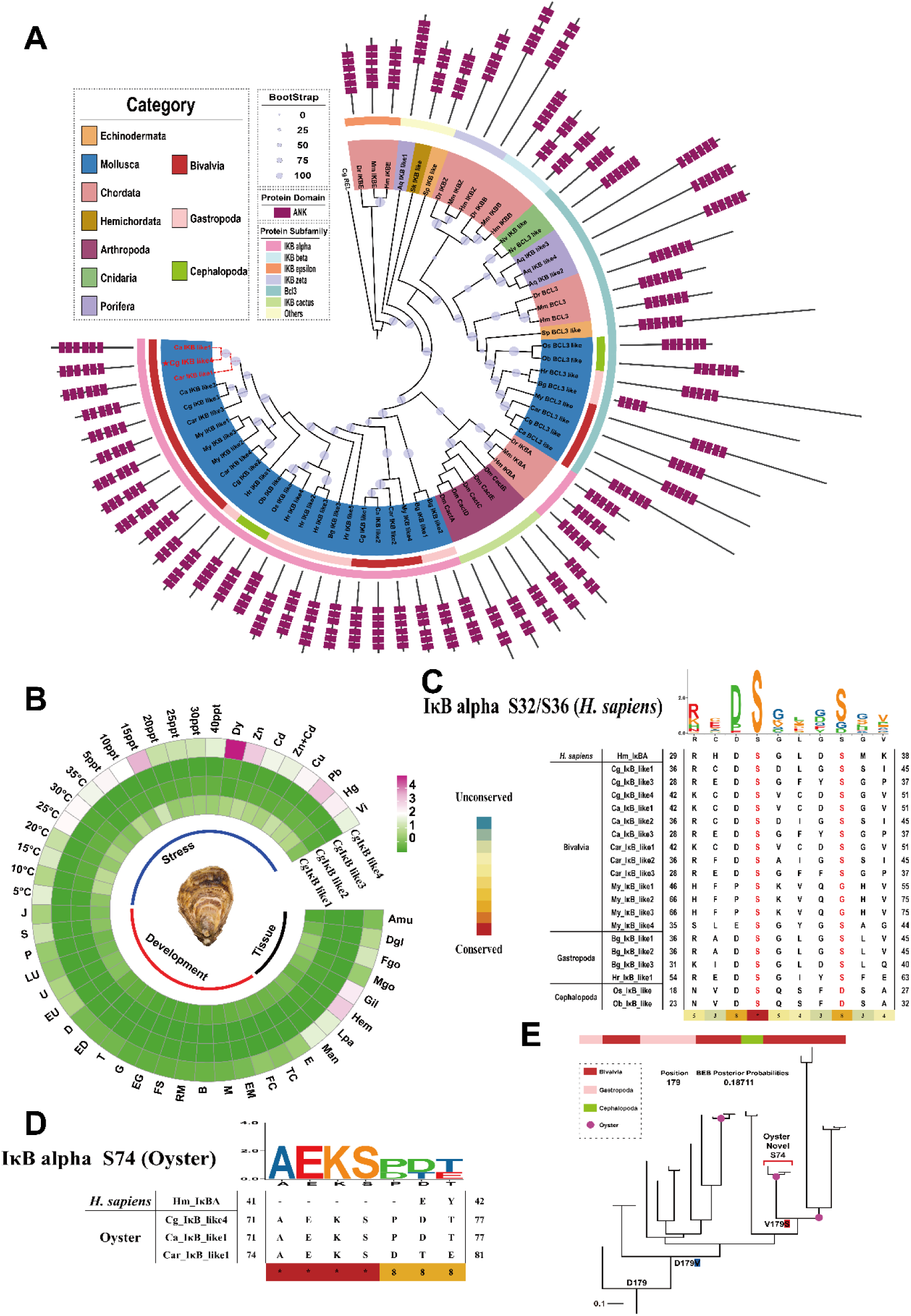
Phylogenetic analysis and sequence alignment of IκB. **A.** Phylogenetic tree of IκB genes family from invertebrates and vertebrates. The tree was constructed with the maximum likelihood (ML) method using PhyloSuite. *Cg*REL was chosen for outgroup. Bootstrap support values are indicated by sizes on nodes of phylogenetic tree. The cluster of *Cg*IκB like4 is marked with red dotted lines. Different colors indicate different categories and IκB subfamilies. Purple rectangle represents ANK domain. Hm, *Homo sapiens*; Mm, *Mus musculus*; Dr, *Danio rerio*; Sk, *Saccoglossus kowalevskii*; Sp, *Strongylocentrotus purpuratus*; Dm, *Drosophila melanogaster*; Cg, *Crassostrea gigas*; Ca, *Crassostrea angulata*; Car, *Crassostrea ariakensis*; My, *Mizuhopecten yessoensis*; Bg, *Biomphalaria glabrata*; Hr, *Haliotis rubra*; Ob, *Octopus bimaculoides*; Os, *Octopus sinensis*; Nv, *Nematostella vectensis*; Aq, *Amphimedon queenslandica*. **B.** Heatmap of four *C. gigas* IκBα genes’ normalized expression (row and column) in different tissues, different developmental stages and under various stress conditions. The relative expression of each gene is indicated by color (from green, low to pink, high). The expression under different stress conditions was all derived from the expression of IκBα in gill tissue. Amu, Adductor; Dgl, Digestive gland; Fgo, Female gonad; Mgo, Male gonad; Gil, Gill; Hem, Hemocyte; Lpa, Labial palp; Man, Mant le; E, egg; TC, Two cells; FC, Four cells; EM, Early morula stage; M, Morula stage; B, Blastula stage; RM, Rotary movement; FS, Free swimming; EG, Early gastrula stage; G, Gastrula stage; T, Trochophore; ED1, Early Dshaped larva; D, Dshaped larva; EU, Early umbo larva; U, Umbo larva; LU, Later umbo larva; P, Competent pediveliger for metamorphosis; S, Spat; J, Juvenile; ppt, Salinity unit (part per thousand); Dy, Drought; Zn, Zinc stress; Cd, Cadmium stress; Zn+Cd, Zinc and Cadmium stress; Cu, Copper stress; Pb, Lead stress; Hg, Mercury stress; Vi, *Vibrio* sp. stress. **C.** The sequence alignment surrounding S32/S36 on human IκBα protein. Different colors correspond to the degree of amino acid conservation (from blue, low to red, high). **D.** The sequence alignment of human and oyster IκBα proteins surrounding S74 on *C. gigas* IκBα like4. **E.** Phylogenetic distribution of mollusca IκBα substitutions at sites 179 (*C. gigas* IκBα like4 S74). Branch lengths represent the number of amino acid substitutions per amino acid site. Bayes empirical Bayes (BEB) posterior probabilities of site 179 under Selection was calculated by PAML.

Furthermore, *Cg* IκB like4 demonstrated responsiveness to abiotic and biotic stressors, including high temperature, salinity, heavy metals, and *Vibrio* sp. stress (Fig. 1B). Additionally, the sequence alignment results showed the presence of conserved Ser32 and Ser36 residues in IκBα proteins of marine mollusks (Fig. 1C). However, the Ser74 residue is unique to oysters (*Cg* IκBα like4, *Ca* IκBα like1, and *Car* IκBα like1), representing specie-specific site (Fig. 1D). To test whether different levels of selective pressures were exerted on oyster’s specific IκBα proteins in mollusks, maximum-likelihood methods implemented in PAML was used to estimate ω value, including specific site and branch models. The likelihood ratio test provided strong evidence favoring the two-ratio model over the one-ratio model, indicating significant heterogeneity in the selective constraint levels between the foreground (oyster-specific IκBα proteins) and background (other mollusks IκBα proteins; P<0.001; Table 1) branches. The oyster-specific IκBα protein (ω_1_=0.00324) exhibited stronger negative selection than other mollusk IκBα proteins (ω_0_=0.18047). Further analysis using site models revealed that under the one-ratio model M0, the maximum likelihood estimate for the ω value of the mollusk IκBα genes was calculated as 0.18, indicating that the expansion of IκBα genes in mollusks was subjected to strong purifying selection (Table 2). Moreover, based on the Δ AIC value, M8a emerged as the most suitable site model, suggesting a certain degree of positive selection acting upon the IκBα gene sequences in mollusks; however, no compelling evidence supports the presence of intense positive selection. The ancestral reconstruction analysis revealed that the amino acid variant V179S (*Cg* IκBα like4, Ser74) was introduced at the onset of the oyster branch, signifying it as a site of oyster-specific evolution (Fig. 1E). The Bayes empirical Bayes posterior probabilities were calculated as 0.1877, suggesting that this site is not subject to positive selection.

**Table 1.**
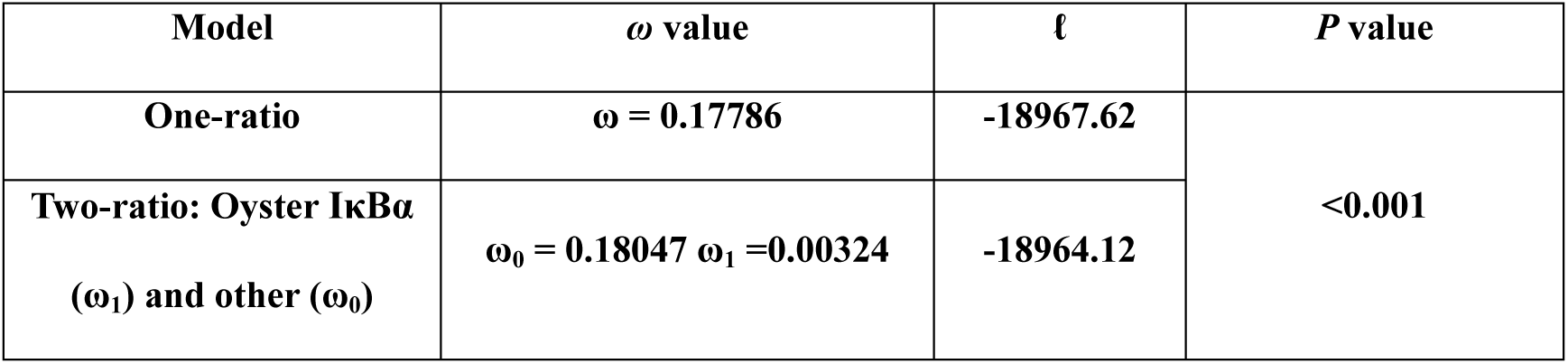
Analyses of strict purifying selection on Oyster-specific IκBα (*Cg*IκBα like4, *Ca*IκBα like1 and *Car*IκBα like1) in Mollusca using PAML branch models.

**Table 2.** Analyses of selection on Oyster-specific IκBα (*Cg*IκBα like4, *Ca*IκBα like1 and *Car*IκBα like1) in Mollusca using PAML random sites models. ^a^ω values of each site class are shown for models M0–M3 (ω_0_–ω_2_) with the proportion of each site class in parentheses. For M7 and M8, the shape parameters, *p* and *q*, which describe the beta distribution are listed instead. In addition, the ω value for the positively selected site class (ω*p*, with the proportion of sites in parentheses) is shown for M8. ^b^Degrees of freedom are given in square brackets after the P-values. ^c^Model fits were assessed by Akaike information criterion differences to the best-fitting model (bolded asterisk). ln *L*, ln Likelihood; N/A, not applicable.

| Model | ln $L$ | Parameters<sup>a</sup> | | | Null | $P$ [df]<sup>b</sup> | $\Delta$ AIC<sup>c</sup> |
| --- | --- | --- | --- | --- | --- | --- | --- |
| | | $\omega_0/p$ | $\omega_1/q$ | $\omega_2/\omega_p$ | | | |
| M0 | -18967.62 | 0.18 | - | - | N/A | - | 1277.04 |
| M1a | -18513.95 | 0.11 (50%) | 1 (50%) | - | M0 | 0.000 [1] | 371.69 |
| M2a | -18513.95 | 0.11 (50%) | 1 (38%) | 1 (11%) | M1a | 1 [2] | 373.69 |
| M3 | -18333.62 | 0.01 (16%) | 0.1 (37%) | 0.42 (47%) | M0 | 0.000 [4] | 15.04 |
| M7 | -18360.65 | 0.59 | 2.28 | - | N/A | - | 83.94 |
| M8a | -18322.28 | 0.68 | 4.45 | 1 | N/A | - | 0<sup>*</sup> |
| M8 | -18324.89 | 0.61 | 2.72 | 6.92 | M7 | 0 [2] | 7.58 |
|  |  |  |  |  | M8a | 0.0222 [1] |  |

### Oyster-specific IκBα inhibited the nuclear entry of the REL to reduce heat resistance

The Co-IP result demonstrated the specific interaction between Myc-*Cg*Rel1, Myc-*Cg*Rel2, and Flag-*Cg*IκB like4 (subsequently referred to as *Cg*IκBα) in the lysates of cells co-transfected with Flag-*Cg*IκBα and Myc-CgRel1 or Myc-CgRel2 (Fig. 2A). Notably, the inability of IgG immunoprecipitation to precipitate REL substantiated the reliability and specificity of the Co-IP results. The BiFC result showed the physical interaction between *Cg*REL1, *Cg*REL2, and *Cg*IκBα in the cytoplasm (Fig. 2B). The subcellular localization results demonstrated that, under normal conditions, the *Cg*REL1 and *Cg*REL2 exhibited a diffuse distribution throughout the cytoplasm and nucleus; however, upon co-transfection with *Cg*IκBα, its localization was prominently constrained to the cytoplasm. High-temperature exposure markedly induced a significant increase in the nuclear translocation of the *Cg*REL1 and *Cg*REL2. However, co-transfection with *Cg*IκBα considerably attenuated the nuclear translocation of REL under a high-temperature condition (Fig. 2C). The luciferase reporter experiments further supported that *Cg*IκBα can gradiently inhibit the positive transcriptional activity of *Cg*REL1 and *Cg*REL2 on the NF-κB signaling pathway promoter (Fig. 2D). The EMSA experiment demonstrated the binding capability of the *Cg*REL1 and *Cg*REL2 to the NF-κB signaling pathway promoter, and the super shift results (Lanes 4 and 8) provided additional evidence confirming its specific binding (Fig. 2E). Western blotting revealed a substantial upregulation of *Cg*REL1 and *Cg*REL2 protein levels in the cellular nucleus under thermal stress, whereas co-transfection with *Cg*IκBα caused a notable decrease in REL nuclear translocation (Fig. 2F). Immunohistochemistry further corroborated the heat-induced nuclear translocation of the REL protein in *C. gigas* and *C. angulata* (Fig. 2G). The cellular heat-induced apoptosis experiments revealed a significant contribution of *Cg*REL1 and *Cg*REL2 in enhancing cellular resistance against high temperature-induced apoptosis (P<0.0001), whereas co-transfection with *Cg*IκBα caused a pronounced REL inhibition, increasing cell apoptosis rate (P<0.0001; Fig. 2H). The CCK-8 cell viability results corroborated the above findings, providing further evidence that the inhibitory effect of *Cg*IκBα on *Cg*REL substantially reduces its contribution to thermotolerance (P<0.001; Fig. 2I). The results of the pilot RNAi experiment indicated that *CgRel1*-1018 (24 h) and *CgRel2*-1813 (24 h) were the most effective siRNA and time for *CgRel1* and *CgRel2*, respectively (Fig. S1). The results of the formal experiment were consistent with those of the pilot experiment, demonstrating the specific knockdown of *CgRel1* and *CgRel2* by *CgRel1*-1018 and *CgRel2*-1813, respectively (Fig. 2J). Following the RNAi experiment, we conducted a heat-induced cell death experiment. The Kaplan–Meier survival analysis demonstrated significantly reduced survival rates in the groups treated with *CgRel1*-1018 and *CgRel2*-1813 (*CgRel1*-siRNA and *CgRel2*-siRNA groups) compared with the control groups (NC and Water groups; P<0.05; Fig. 2K).

**Fig. 2.**
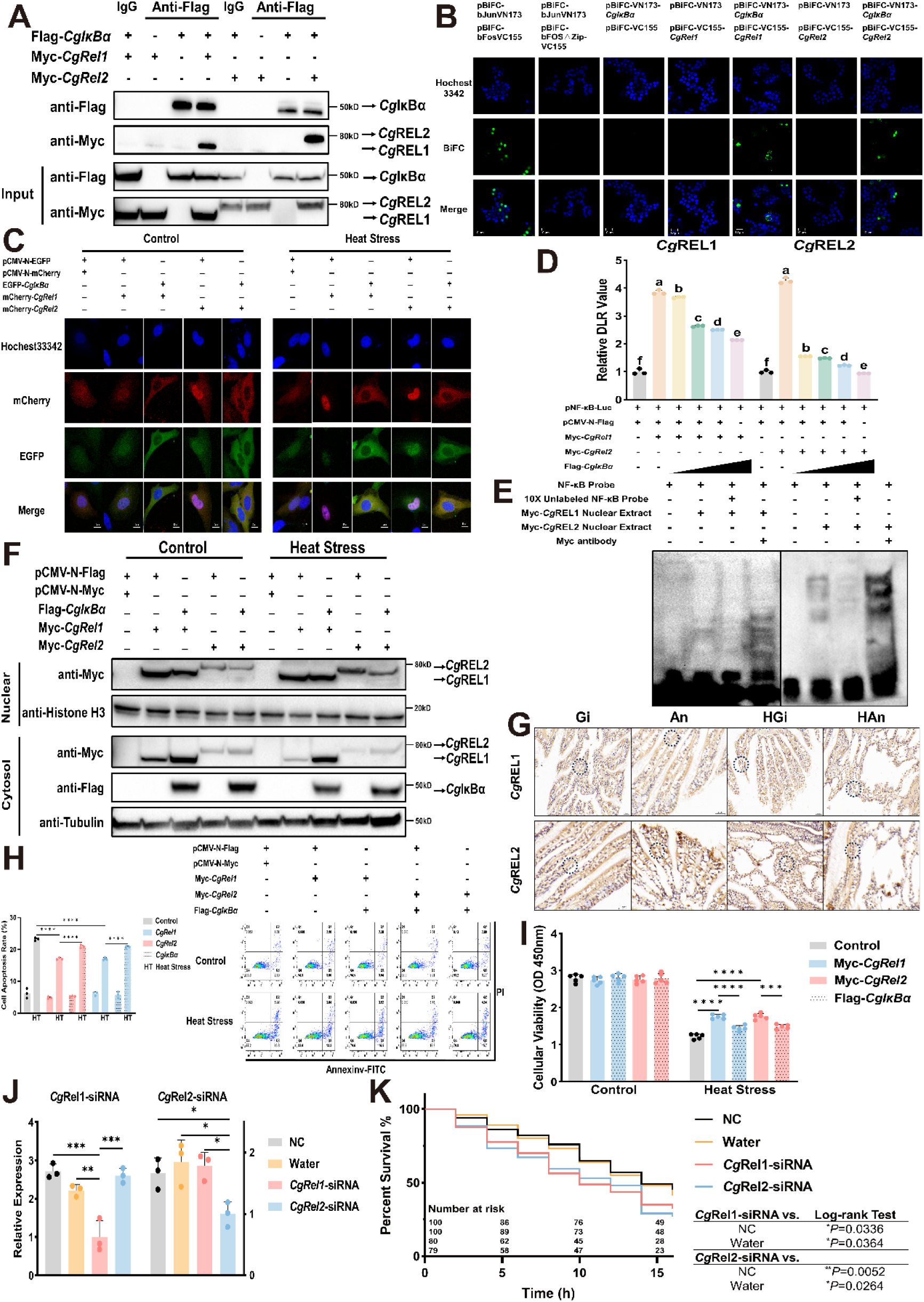
*Cg*IκBα inhibits the nuclear translocation of *Cg*REL and then decreases its heat resistance. **A.** Co-immunoprecipitation (co-IP) of *Cg*IκBα with *Cg*REL1/*Cg*REL2. HEK293T cells expressing the indicated constructs encoding Flag-*CgIκBα* and Myc-*CgRel1*/Myc-*CgRel2* were lysed and incubated with anti-Flag magnetic beads overnight, Myc-tagged molecules co-IPed in this manner were resolved by SDS-PAGE and detected by immunoblotting with anti-Myc antibody. The expression of *Cg*IκBα and *Cg*REL1/*Cg*REL2 by transfectants (INPUT) in these studies was also confirmed by immunoblot analysis. **B.** BiFC assay of *Cg*IκBα and *Cg*REL1/*Cg*REL2. HeLa cells were transfected BiFC plasmids expressing *Cg*IκBα (pBiFC-VN173-*CgIκBα*) only, *Cg*REL1 (pBiFC-VC155-*CgRel1*)/*Cg*REL2 (pBiFC-VC155-*CgRel2*) only or *Cg*IκBα and *Cg*REL1/*Cg*REL2. Images were acquired with a confocal microscope at the EGFP channel. Bar: 10 µm. **C.** Subcellular localization of *Cg*IκBα and *Cg*REL1/*Cg*REL2 in HeLa cells under control and heat stress. HeLa cells were transfected mCherry-*CgRel1*/mCherry*CgRel2* only, or EGFP-*CgIκBα* and mCherry-*CgRel1*/mCherry-*CgRel2*. Images were acquired with a confocal microscope under control and heat treatment. Bar: 10 µm. **D.** The relative dual-luciferase reporter (DLR) values of HEK293T cells transfected with pNF-κB-Luc, pRL-TK, Myc-*CgRel1*/Myc-*CgRel2* and Flag-*CgIκBα* (n=3). Among these, the transfection amounts of Flag-*CgIκBα* were increased in a gradient manner, with doses of 100 ng, 300 ng, 500 ng, and 700 ng per well of a 24-well plate, respectively. The error bars represent the S.D. **E.** EMSA assay of the biotin-labeled NF-κB consensus oligo with *Cg*REL1/*Cg*REL2 transfected cell nuclear extract. The unlabeled probes added at 10-fold excess were used to verify specific DNA–protein interactions (lines 3, 7). And the 1 µg Myc-antibody was added in the reaction mixture for super-shift experiment. **F.** The western blotting of nuclear and cytoplasmic proteins from HEK292T cells transfected with Flag-*CgIκBα* and Myc-*CgRel1*/Myc-*CgRel2* under control and heat treatment. **G.** The immunohistochemistry of *Cg*REL1 and *Cg*REL2 in gill tissues from *C. gigas* and *C. angulata* under control and heat treatment. The cell apoptosis rate (**H.**; n=3) and the CCK-8 assay for the cell viability (**I.**; n=5) of HEK293T cells transfected with Flag-*CgIκBα* and Myc-*CgRel1*/Myc-*CgRel2* under control and heat treatment. The left panel is the cell apoptosis chart in each group, and the right panel is the cell apoptosis rate in each group. The error bars represent the S.D. **J.** The relative expression of oyster *Rel1* and *Rel2* in gill tissues in *Cg*REL RNA interference experiments (n=3). NC, Water, *Rel1*-siRNA and *Rel2*-siRNA in legend represent the oysters were injected with nonsense strands, water, siRNA, respectively. The error bars represent the S.D. **K.** Kaplan-Meier survival curves of oysters in heat-lethal experiments after *Cg*REL RNA interference experiments. Different colors represent oysters injected with nonsense strands, water, siRNA, respectively. Significant differences among groups were marked with \**p*<0.05, \*\**p*<0.01, \*\*\**p*<0.001, and \*\*\*\**p*<0.0001. “ns” indicates non-significant differences.

### Phosphorylation at Ser74 independently mediated the ubiquitination and degradation of the oyster-specific IκBα

Our previous study found divergent phosphorylation levels at the Ser74 site of the oyster-specific IκBα (*Cg*IκB like4) during heat stress in *C. gigas* and *C. angulata*. *Crassostrea angulata* exhibited a significant upregulation of Ser74 phosphorylation (P<0.01), whereas no detectable phosphorylation at S74 was observed in *C. gigas* after heat stress (Fig. 3A). The site mutation protein degradation experiments demonstrated substantial inhibition of ubiquitination and degradation of the *Cg*IκBα protein when treated with a combination of MG132 and CHX. When treated with CHX alone, the *Cg*IκBα^S74D^ mutant (mimicking phosphorylation)/*Cg*IκBα^S74A^ mutant (mimicking dephosphorylation) exerted an augmenting/inhibitory effect on *C*gIκBα degradation.

**Fig. 3.**
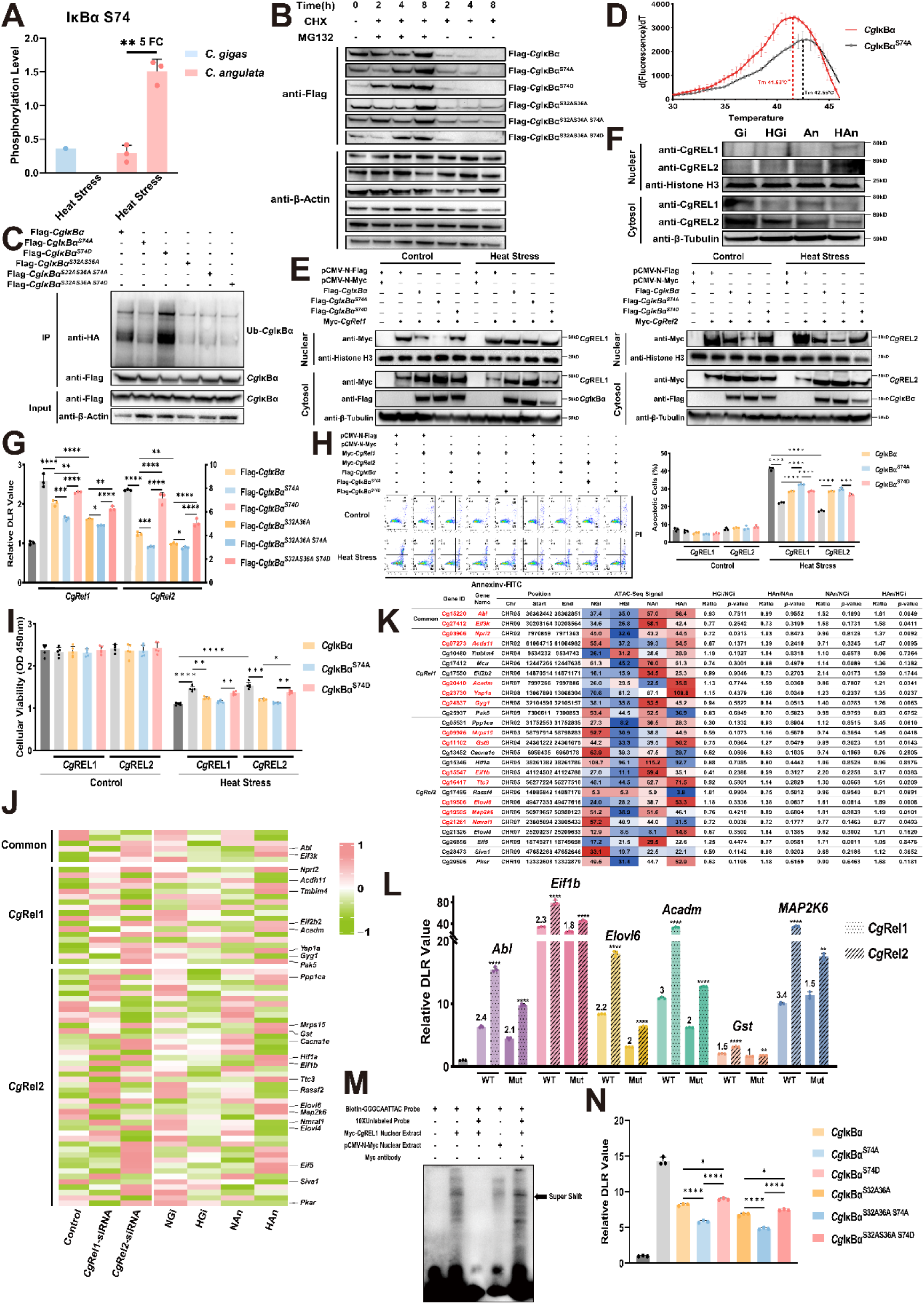
The function of phosphorylation at Ser74 of *Cg*IκBα and downstream genes regulated by *Cg*REL. **A.** The phosphorylation levels of Ser74 site of *Cg*IκBα in gill tissues of *C. gigas* and *C. angulata* during heat stress, which was obtained from our previous study (Wang et al., 2023a). **B.** The protein degradation experiments of different *Cg*IκBα mutants. The HEK293T cells were transfected with Flag-*CgIκBα*, Flag-*CgIκBα^S74A^*, Flag-*CgIκBα^S74D^*, Flag-*CgIκBα^S32AS36A^*, Flag-*CgIκBα^S32AS36A^ ^S74A^*, and Flag-*CgIκBα^S32AS36A^ ^S74D^* and incubated with CHX (50 μM) only, and MG132 (10 μM) and CHX in combination for 2, 4, 8 hours. The content of different *Cg*IκBα mutants was measured by immunoblot analysis anti-Flag antibody. **C.** The protein ubiquitination experiments of different *Cg*IκBα mutants. The HEK293T cells were co-transfected with HA-Ub and Flag-*CgIκBα*/Flag-*CgIκBα^S74A^*/Flag-*CgIκBα^S74D^*/Flag-*CgIκBα^S32AS36A^*/Flag-*CgIκBα^S32AS3A6^ ^S74A^*/Flag-*CgIκBα^S32AS36A^ ^S74D^*, and incubated with 10 μM MG132 for 8 hours. Then, the cells were lysed and incubated with anti-Flag magnetic beads overnight, HA-tagged molecules co-IPed in this manner were resolved by SDS-PAGE and detected by immunoblotting with anti-HA antibody, which used to measure the ubiquitination level of different *Cg*IκBα mutants. **D.** Thermal stabilization of *Cg*IκBα and *Cg*IκBα^S74A^ proteins, as measured via differential scanning fluorimetry (DSF). Mean Tm values ± S.D. is shown (n = 3). \**p* < 0.05 by two-tailed unpaired t test. **E.** The western blotting of nuclear and cytoplasmic proteins from HEK292T cells transfected with Flag-*CgIκBα*/Flag-*CgIκBα^S74A^*/Flag-*CgIκBα^S74D^*and Myc-*CgRel1*/Myc-*CgRel2* under control and heat treatment. **F.** The western blotting of nuclear and cytoplasmic proteins extracted from gill tissues of *C. gigas* and *C. angulata* during heat stress. The content of *Cg*REL1 and *Cg*REL2 were detected by immunoblotting with anti-*Cg*REl1/*Cg*REL2 antibody. Gi and An represent the *C. gigas* and *C. angulata* under control condition. HGi and HAn represent the *C. gigas* and *C. angulata* under heat stress. **G.** The relative dual-luciferase reporter (DLR) values of HEK293T cells transfected with pNF-κB-Luc, pRL-TK, Myc-*CgRel1*/Myc-*CgRel2* and Flag-*CgIκBα*/Flag-*CgIκBα^S74A^*/Flag-*CgIκBα^S74D^*/Flag-*CgIκBα^S32AS36A^*/Flag-*CgIκBα^S32AS3A6^ ^S74A^*/Flag-*CgIκBα^S32AS36AS74D^* (n=3). The error bars represent the S.D. The cell apoptosis rate (**H.**; n=3) and the CCK-8 assay for the cell viability (**I.**; n=5) of HEK293T cells transfected with Flag-*CgIκBα*/Flag-*CgIκBα^S74A^*/Flag-*CgIκBα^S74D^*and Myc-*CgRel1*/Myc-*CgRel2* under control and heat treatment (n=3). The left panel is the cell apoptosis chart in each group, and the right panel is the cell apoptosis rate in each group. The error bars represent the S.D. **J.** The heatmap of *Cg*REL-regulated downstream genes normalized expression (row) from transcriptomic data of RNAi experiment and *C. gigas* and *C. angulata* during heat stress (Wang et al., 2024). The relative expression of each gene is indicated by color (from green, low to pink, high). **K.** The values of ATAC-Seq signal in the *Cg*REL binding regions within promoter of *Cg*REL-regulated downstream genes from *C. gigas* and *C. angulata* under heat stress (Wang et al., 2024). **L.** The relative dual-luciferase reporter (DLR) values of cells co-transfected with the promoter of the *Abl*/*Eif1b*/*Elovl6*/*Acadm*/*Gst*/*Map2k6* or their corresponding *Cg*REL binding region deletion mutations ligated with pGL3-basic plasmid, and Myc-*CgRel1*/Myc-*CgRel2* (n=3). The error bars represent the S.D. **M.** EMSA assay of the biotin-labeled REL-binding sequences in the promoter of *Acadm*, which was predicted by the JASPAR database, with *Cg*REL1 transfected cell nuclear extract. The unlabeled probes added at 10-fold excess were used to verify specific DNA–protein interactions (line 3). The nuclear extract from cell transfected with empty plasmid (pCMV-N-Myc) was used to demonstrate the HEK293T cells endogenous binding (line 4). And the 1 µg Myc-antibody was added in the reaction mixture for super-shift experiment (line 5). **N.** The relative dual-luciferase reporter (DLR) values of cells co-transfected with the promoter of the *Acadm* or their corresponding *Cg*REL binding region deletion mutation ligated with pGL3-basic plasmid, Myc-*CgRel1*, and Flag-*CgIκBα*/Flag-*CgIκBα^S74A^*/Flag-*CgIκBα^S74D^*/Flag-*CgIκBα^S32AS36A^*/Flag-*CgIκBα^S32AS3A6^ ^S74A^*/Flag-*CgIκBα^S32AS36A^ ^S74D^* (n=3). The error bars represent the S.D. Significant differences among groups were marked with \**p*<0.05, \*\**p*<0.01, \*\*\**p*<0.001, and \*\*\*\**p*<0.0001. “ns” indicates non-significant differences.

This effect was observed to be independent of phosphorylation at S32 and S36 (Fig. 3B). Furthermore, the protein ubiquitination experiments revealed results consistent with the aforementioned findings, demonstrating that phosphorylation at S74 significantly enhanced the ubiquitination level of the *Cg*IκBα protein (Fig. 3C). Interestingly, the DSF results demonstrated that the simulation of dephosphorylation at S74 significantly increased the thermal stability of the *Cg*IκBα protein (P<0.05; Fig. 3D). Western blotting showed that phosphorylation at the S74 can enhance the nuclear translocation of *Cg*REL1 and *Cg*REL2 by facilitating *Cg*IκBα degradation (Fig. 3E). Moreover, the nuclear and cytoplasmic Western blotting on *C. gigas* and *C. angulata* after heat stress significantly supported the above results. *Crassostrea angulata* with high S74 phosphorylation levels exhibited a higher abundance of nuclear REL protein after heat stress (Fig. 3F). The luciferase reporter gene experiments further confirmed that phosphorylation of *Cg*IκBα at S74 promoted its degradation independent of S32 and S36, relieving the inhibition of *Cg*REL and enhancing its transcriptional activity in activating the NF-κB signaling pathway promoter (Fig. 3G). In addition, the cellular heat-induced apoptosis experiments (P<0.001; Fig. 3H) and CCK-8 cell viability assays (P<0.01; Fig. 3I) indicated that phosphorylation of *Cg*IκBα at the S74 site enhanced the role of *Cg*REL in thermotolerance by promoting its ubiquitin-mediated degradation.

### Differential phosphorylation of *Cg*IκBα^S74^ activated cell survival, fatty acid metabolism, translation, and antioxidant-related genes to mediate divergent heat response

DAP-Seq was performed to obtain the downstream genes regulated by *Cg*REL1 and *Cg*REL2. A total of 2.68 × 10^10^ clean reads were first obtained, and 78% of reads were mapped to the *C. gigas* genome. The overall information on alignment is shown in Figure S2. Based on peak calling results from MACS2 software, 6,208 and 8,195 peaks (q<0.05) were identified in *Cg*REL1 and *Cg*REL2, with an average length of 482 and 482 bp, respectively. The peak distribution demonstrated that the binding regions of *Cg*REL1 and *Cg*REL2 were primarily concentrated within a 2-kb interval upstream and downstream of the genes (Fig. S3A, D). At the chromosomal level, chr5, chr2, chr9, and chr6 were identified as the top four chromosomes with the highest peak enrichment in *Cg*REL1, with 824, 803, 763, and 758 peaks, respectively (Fig. S3B). In *Cg*REL2, chr5, chr2, chr6, and chr7 were the top four chromosomes, with 1,162, 1,117, 1,014, and 947 peaks, respectively (Fig. S3E). Genomic region analysis of *Cg*REL1/*Cg*REL2 showed that 25.51/21.63%, 20.92/21.65%, and 7.86/8.76% of the peaks were in the intron, distal intergenic region, and promoter region, respectively (Fig. S3C, F). We screened the peaks localized within the promoter regions of genes and identified 349, 570, and 135 potential genes regulated by *Cg*REL1, *Cg*REL2, and *Cg*REL1 and *Cg*REL2, respectively (Table S5). The transcriptomic data following the RNAi experiment showed an average of 30,633,206 high-quality 150-bp paired-end reads mapped to the *C. gigas* genome, with mapping rates of 71.83% (Table S6). Principal component analysis was performed to compare the transcriptomic data of the three groups, which showed that PC1 and PC2 accounted for 32.7% and 26.1% of the total variance, respectively. The three groups exhibited complete separation, indicating the robustness of the RNAi knockdown (Fig. S4; Table S7). By integrating DAP-Seq and transcriptomic data after the RNAi experiment, along with existing gene function studies, we manually identified 69 potential target genes regulated by *Cg*REL involved in biological temperature adaptation (Fig. 3J). These genes exhibited DAP-Seq peaks within their promoter regions and demonstrated differential expression levels between *Cg*Rel1-siRNA/*Cg*Rel2-siRNA and control groups (Table S8). Furthermore, by incorporating previous transcriptomic data from *C. gigas* and *C. angulata* under heat stress, 26 genes were identified as differentially activated during the heat stress responses between these species (Table S8). Additionally, using previous ATAC-Seq data from *C. gigas* and *C. angulata* under heat stress, ATAC-Seq peak signals were collected from the *Cg*REL binding regions in the promoters of the above 26 genes. The results showed that 14 genes were identified with significantly higher ATAC-Seq signals in heat-stressed *C. angulata* than in *C. gigas* (Fig. 3K). Among them, two (including tyrosine-protein kinase Abl (*Abl*)) were co-regulated by *Cg*REL1 and *Cg*REL2, whereas five (such as medium-chain specific acyl-CoA dehydrogenase, mitochondrial (*Acadm*)) were specifically regulated by *Cg*REL1 and seven were specifically regulated by *Cg*REL2 (such as glutathione S-transferase 8 (*Gst*)). By combining the binding region deletion mutations and luciferase reporter assays, we successfully identified two (*Abl* and *Acadm*) and four (eukaryotic translation initiation factor 1 (*Eif1b*), elongation of very long chain fatty acids protein 6 (*Elovl6*), dual specificity mitogen-activated protein kinase kinase 6 (*Map2k6*), and *Gst*) genes specifically activated by *Cg*REL1 and *Cg*REL2, respectively, which exhibited lower increase in fluorescence intensity in the mutant-type transfected with *Cg*REL than the wild-type (Fig. 3L). Using *Acadm* as an example, the REL1 binding region within its promoter region exhibited differential accessibility during heat stress in *C. gigas* and *C. angulata* (Fig. S5), suggesting that *Cg*REL1 may enhance the expression of *Acadm* in *C. angulata* under heat stress by increasing its interactions with the binding region of the promoter. Moreover, the EMSA experiment was performed to confirm the specificity of CgREL1 binding to the promoter region of *Acadm*. The transfection of Myc-*CgRel1* into cellular nuclear extract caused significant retardation of band migration (lanes 1, 2, 3, 4), and the results of super shift further supported the binding specificity (Lane 5; Fig. 3M). Additionally, we performed repeated validation of the ability of *Cg*IκBα S74 phosphorylation to independently mediate protein degradation using the luciferase reporter assay and *Acadm* promoter (Fig. 3N).

### *Cg*ERK1/2 phosphorylated *Cg*IκBα S74 to promote its ubiquitin-mediated protein degradation

Based on the results of kinase prediction, we identified numerous kinases associated with the MAPK pathway, with a particular emphasis on the ERK genes (Table S9). Phylogenetic analysis confirmed the identification of oyster ERK protein as ERK1/2 (Fig. 4A). The Co-IP result demonstrated the specific interaction between *Cg*ERK1/2 and *Cg*AKT in the lysates of cells co-transfected with Flag-*CgIκBα* and Myc-*CgErk1/2* (Fig. 4B). The yeast two-hybrid (Fig. 4C) and BiFC assays (Fig. 4D) provided additional evidence supporting the specific interaction between *Cg*ERK1/2 and *Cg*IκBα, predominantly localized within the cytoplasm. The subcellular localization results demonstrated that *Cg*ERK1/2 markedly attenuated the inhibitory effect of *Cg*IκBα on *Cg*REL, consequently promoting nuclear translocation of REL proteins in response to heat stress (Fig. 4E). The *in vivo* kinase experiment indicated that *Cg*ERK1/2 phosphorylated the S74 site of *Cg*IκBα, demonstrated by the more pronounced migration bands observed when co-transfected with *Cg*ERK1/2 and *Cg*IκBα^S32AS36A^ than that with Lane 1 that transfected with *Cg*IκBα^S32AS36A^ alone (Fig. 4F). The migration bands of *Cg*IκBα^S32AS36A^ ^S74A^ mutant showed no significant changes regardless of co-transfection with *Cg*ERK1/2. Moreover, the *in vitro* kinase experiments further support the above findings. Co-incubation of *Cg*ERK1/2 with *Cg*IκBα resulted in a noticeable band migration, eliminated upon co-incubation of *Cg*IκBα^S74A^ and *Cg*ERK1/2 (Fig. 4G). Furthermore, the protein ubiquitination experiment showed that co-transfection of *Cg*ERK1/2 and *Cg*IκBα^S32AS36A^ caused a significant increase in the ubiquitination level of *Cg*IκBα, which was effectively weakened by the *Cg*IκBα^S32AS36A^ ^S74A^ mutant (Fig. 4H). The protein degradation experiments demonstrated that *Cg*ERK1/2 significantly promoted *Cg*IκBα degradation, which was unaffected by phosphorylation at S32 and S36. The S74 dephosphorylation mutation effectively prevented the *Cg*IκBα protein degradation by blocking the ERK-mediated phosphorylation at S74 (Fig. 4I). Western blotting on heat-stress cells further confirmed that *Cg*ERK1/2 facilitated the degradation of *Cg*IκBα, enabling increased nuclear translocation of *Cg*REL1 and *Cg*REL2 during high temperatures, which was inhibited by the S74 dephosphorylation mutation (Fig. 4J). Additionally, the luciferase reporter experiments demonstrated that *Cg*ERK1/2, independent of phosphorylation at S32 and S36, can gradiently increase the activation of *Cg*REL-mediated NF-κB signaling pathway transcriptional activity (Fig. 4K). The cellular heat-induced apoptosis experiments (Fig. 4L) and the corresponding CKK-8 cell viability measurements (Fig. 4M) provided additional evidence that *Cg*ERK1/2 is crucial in mitigating the inhibitory effect of *Cg*IκBα on *Cg*REL1 and *Cg*REL2 and that it enhances *Cg*REL’s role in facilitating cell resistance against high-temperature-induced cell death.

**Fig. 4.**
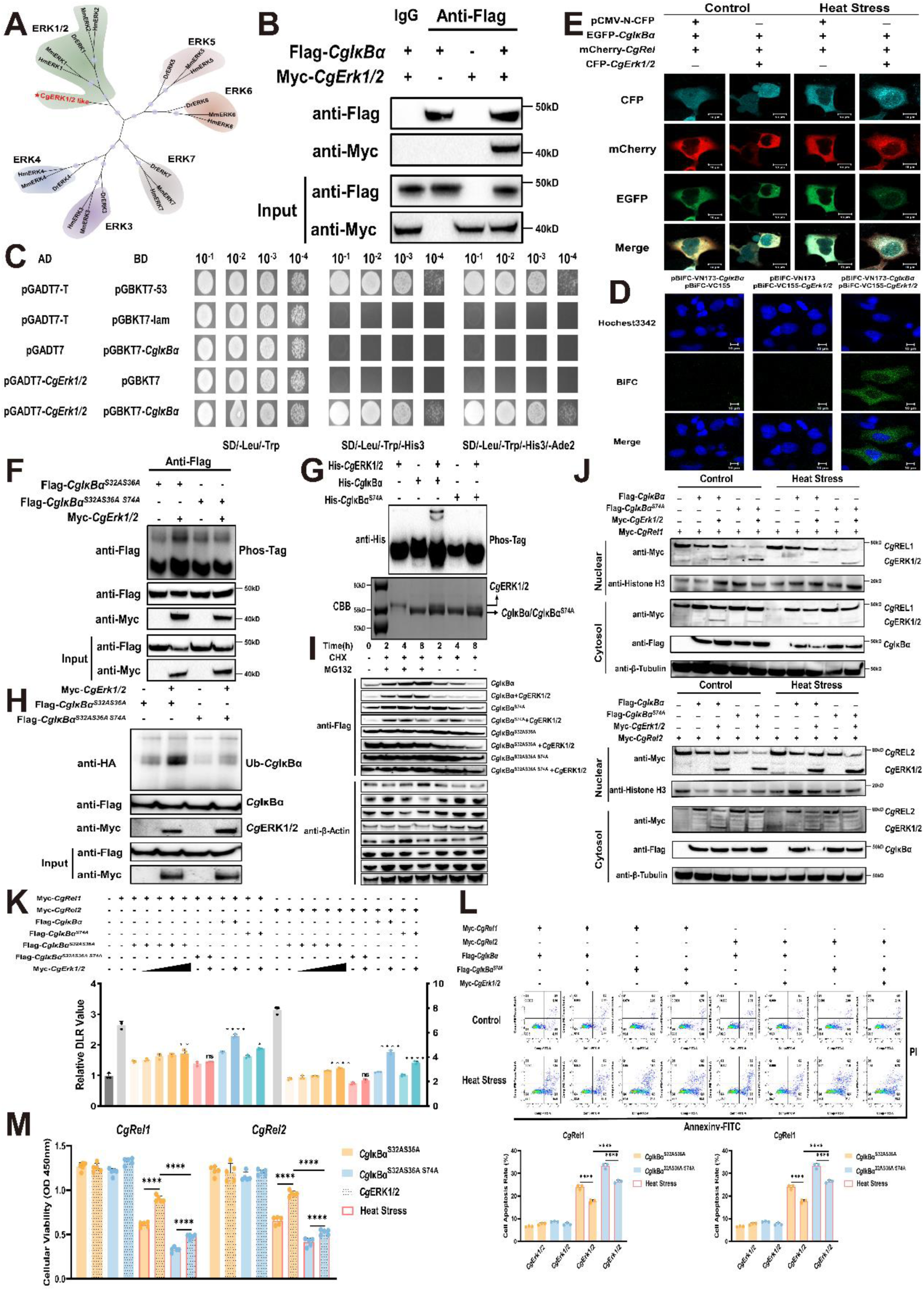
The interaction and kinase reaction between *Cg*ERK1/2 and *Cg*IκBα. **A.** Phylogenetic tree of oyster, human, mouse and zebrafish ERK gene family. The tree was constructed with the maximum likelihood (ML) method using PhyloSuite. Bootstrap support values are indicated by sizes on nodes of phylogenetic tree. The *Cg*ERK1/2 is marked with red bold line. Different colors indicate different categories of ERK subfamilies. Cg, *Crassostrea gigas*; Hm, *Homo sapiens*; Mm, *Mus musculus*; Dr, *Danio rerio*. The accession numbers were as follows: *Hm* ERK1, NP_002737.2; *Hm* ERK2, NP_002736.3; *Hm* ERK3, NP_002739.1; *Hm* ERK4, NP_002738.2; *Hm* ERK5, NP_002740.2; *Hm* ERK6, NP_002960.2; *Hm* ERK7, NP_620590.2; *Mm* ERK1, NP_036082.1; *Mm* ERK2, NP_001033752.1; *Mm* ERK3, NP_056621.4; *Mm* ERK4, NP_766220.2; *Mm* ERK5, NP_001277963.1; *Mm* ERK6, NP_001389948.1; *Mm* ERK7, NP_808590.1; *Dr* ERK1, NP_958915.1; *Dr* ERK3, NP_001039017.1; *Dr* ERK4, NP_998638.1; *Dr* ERK5, NP_001013469.2; *Dr* ERK6, NP_571482.1; *Dr* ERK7, XP_009292731.1. The sequences of *Cg*ERK1/2 (Cg03498) gene in oyster were obtained from *Crassostrea gigas* genome blast (GCA_011032805.1). **B.** Co-immunoprecipitation (co-IP) of *Cg*IκBα with *Cg*ERK1/2. HEK293T cells expressing the indicated constructs encoding Flag-*CgIκBα* and Myc-*CgErk1/2* were lysed and incubated with anti-Flag magnetic beads overnight, Myc-tagged molecules co-IPed in this manner were resolved by SDS-PAGE and detected by immunoblotting with anti-Myc antibody. The expression of *Cg*IκBα and *Cg*ERK1/2 by transfectants (INPUT) in these studies was also confirmed by immunoblot analysis. **C.** Yeast two-hybrid assay between *Cg*IκBα and *Cg*ERK1/2. Full-length of *CgIκBα* and *CgErk1/2* were fused to the pGBKT7 binding domain (BD, bait) and the pGADT7 activation domain (AD, prey), respectively, and then transformed into yeast. Shown are growth phenotypes of yeast transformants on selective media of SD/Leu-Trp-(left panel), SD/Leu-Trp-His-(central panel, interaction) and SD/Leu-Trp-His-Ade2 (right panel, interaction). **D.** BiFC assay of *Cg*IκBα and *Cg*ERK1/2. HeLa cells were transfected BiFC plasmids expressing *Cg*IκBα (pBiFC-VN173-*CgIκBα*) only, *Cg*ERK1/2 (pBiFC-VC155-*CgErk1/2*) only or *Cg*IκBα and *Cg*ERK1/2. Images were acquired with a confocal microscope at the EGFP channel. Bar: 10 µm. **E.** Subcellular localization of *Cg*IκBα, *Cg*REL and *Cg*ERK1/2 in HeLa cells under control and heat stress. HeLa cells were transfected mCherry-*CgRel* and EGFP-*CgIκBα* or mCherry-*CgRel*, EGFP-*CgIκBα* and CFP-*CgErk1/2*. Images were acquired with a confocal microscope under control and heat treatment. Bar: 10 µm. **F.** *In vivo* phosphorylation assay of *Cg*ERK1/2 on *Cg*IκBα Ser74 site. HEK293T cells were transfected with Flag-*CgIκBα^S32AS36A^*/Flag-*CgIκBα^S32AS36A^ ^S74A^* and Myc-Cg*Erk1/2*. The cells were lysed and incubated with anti-Flag magnetic beads overnight, and IPed proteins were separated using SDS-PAGE gels containing Phos-tag (+Phos-tag). The phosphorylation level of Flag-*Cg*IκBα^S32AS36A^/Flag-*Cg*IκBα^S32AS36A^ ^S74A^ were detected by western blotting using anti-Flag antibody. Protein samples were also separated in SDS-PAGE gels without Phos-tag (–Phos-tag). **G.** *In vitro* kinase activity assay of *Cg*ERK1/2 on *Cg*IκBα Ser74 site. The recombinant proteins (His-*Cg*ERK1/2 and His-*Cg*IκBα/His-*Cg*IκBα^S74A^) were incubated in kinase buffer at 30 °C for 30 min and separated using SDS-PAGE gels containing Phos-tag, which were detected by western blotting with anti-His antibody. CBB, Coomassie Brilliant Blue staining. **H.** The protein ubiquitination experiments of *Cg*IκBα^S32AS36A^/*Cg*IκBα^S32AS36A^ ^S74A^ co-transfected with CgERK1/2. The HEK293T cells were co-transfected with HA-Ub, Flag-*CgIκBα^S32AS36A^*/Flag-*CgIκBα^S32AS36A^ ^S74A^* and Myc-*CgErk1/2*, and incubated with 10 μM MG132 for 8 hours. Then, the cells were lysed and incubated with anti-Flag magnetic beads overnight, HA-tagged molecules co-IPed in this manner were resolved by SDS-PAGE and detected by immunoblotting with anti-HA antibody, which used to measure the ubiquitination level of *Cg*IκBα^S32AS36A^ and *Cg*IκBα^S32AS36A^ ^S74A^. **I.** The protein degradation experiments of *Cg*IκBα/*Cg*IκBα^S74A^/*Cg*IκBα^S32AS36A^/*Cg*IκBα^S32AS36A^ ^S74A^ co-transfected with CgERK1/2. The HEK293T cells were transfected with Flag-*CgIκBα*, Flag-*CgIκBα^S74A^*, Flag-*CgIκBα^S32AS36A^*, Flag-*CgIκBα^S32AS36A^ ^S74A^* only, or co-transfected with Myc-*CgErk1/2*, and incubated with CHX (50 μM) only, and MG132 (10 μM) and CHX in combination for 2, 4, 8 hours. The content of *Cg*IκBα in each group was measured by immunoblot analysis anti-Flag antibody. **J.** The western blotting of nuclear and cytoplasmic proteins from HEK292T cells transfected with Flag-*CgIκBα*/Flag-*CgIκBα^S74A^*, Myc-*CgRel1*/Myc-*CgRel2* and Myc-*CgErk1/2* under control and heat treatment. **K.** The relative dual-luciferase reporter (DLR) values of HEK293T cells transfected with pNF-κB-Luc, pRL-TK, Myc-*CgRel1*/Myc-*CgRel2*, Flag-*CgIκBα*/Flag-*CgIκBα^S74A^*/Flag-*CgIκBα^S32AS36A^*/Flag-*CgIκBα^S32AS36A^ ^S74A^* and Myc-*CgErk1/2* (n=3). Among these, the transfection amounts of Myc-*CgErk1/2* was increased in a gradient manner, with doses of 100 ng, 200 ng, 300 ng, and 400 ng per well of a 24-well plate, respectively. The error bars represent the S.D. The cell apoptosis rate (**L.**; n=3) and the CCK-8 assay for the cell viability (**M.**; n=5) of HEK293T cells transfected with Flag-*CgIκBα^S32AS36A^*/Flag-*CgIκBα^S32AS36A^ ^S74A^*, Myc-*CgRel1*/Myc-*CgRel2* and Myc-*CgErk1/2* under control and heat treatment (n=3). The upper panel is the cell apoptosis chart in each group, and the lower panel is the cell apoptosis rate in each group. The error bars represent the S.D. Significant differences among groups were marked with \**p*<0.05, \*\**p*<0.01, \*\*\**p*<0.001, and \*\*\*\**p*<0.0001. “ns” indicates non-significant differences.

### Divergent phosphorylation of *Cg*ERK1/2^T187^ ^Y189^ by MAPK signaling pathway mediated differential NF-κB pathway activity

The sequence alignment results showed that the oyster ERK1/2 residues T187 and Y189 correspond to the conserved active sites T202 and Y204, respectively, in human ERK1 (Fig. 5A). Previous phosphoproteomic data from *C. gigas* and *C. angulata* during heat stress revealed distinct patterns of phosphorylation at the *Cg*ERK1/2 T187 and Y189 sites. The phosphorylation level at the *Cg*ERK1/2 T187 site increased in *C. angulata* and decreased in *C. gigas* and significantly increased at the *Cg*ERK1/2 Y189 site only in *C. angulata* under heat stress (P<0.05; Fig. 5B). *In vivo* (Fig. 5C) and *in vitro* (Fig. 5D) kinase assays, coupled with *Cg*ERK1/2^T187D^ ^Y189E^ (mimicking phosphorylation) and *Cg*ERK1/2^T187A^ ^Y189F^ (mimicking dephosphorylation) mutants provided compelling evidence that phosphorylation at *Cg*ERK1/2 T187 and Y189 significantly enhanced its kinase activity, causing an elevated phosphorylation level of *Cg*IκBα S74 site. In addition, the protein ubiquitination experiments showed that phosphorylation at *Cg*ERK1/2 T187 and Y189 significantly enhanced the ubiquitination level of *Cg*IκBα^S32A^ ^S36A^ (Fig. 5E). Western blotting on heat-stress cells further demonstrated that phosphorylation at *Cg*ERK1/2 T187 and Y189 contributed to the enhancement of its kinase activity, promoting *Cg*IκBα degradation to facilitate the nuclear translocation of *Cg*REL1 and *Cg*REL2 in response to heat stress (Fig. 5F). Similarly, the *Cg*ERK1/2 T187 and Y189 phosphorylation mutants significantly enhanced REL-mediated positive transcriptional activity in the NF-κB signaling pathway (P<0.05; Fig. 5G). Furthermore, the cellular heat-induced apoptosis experiments (Fig. 5H) and CCK-8 cell viability measurements (Fig. 5I) provide additional evidence supporting the role of *Cg*ERK1/2 T187 and Y189 phosphorylation in facilitating *Cg*IκBα^S32AS36A^ degradation and amplifying the protective effect of *Cg*REL against heat stress-induced cell death.

**Fig. 5.**
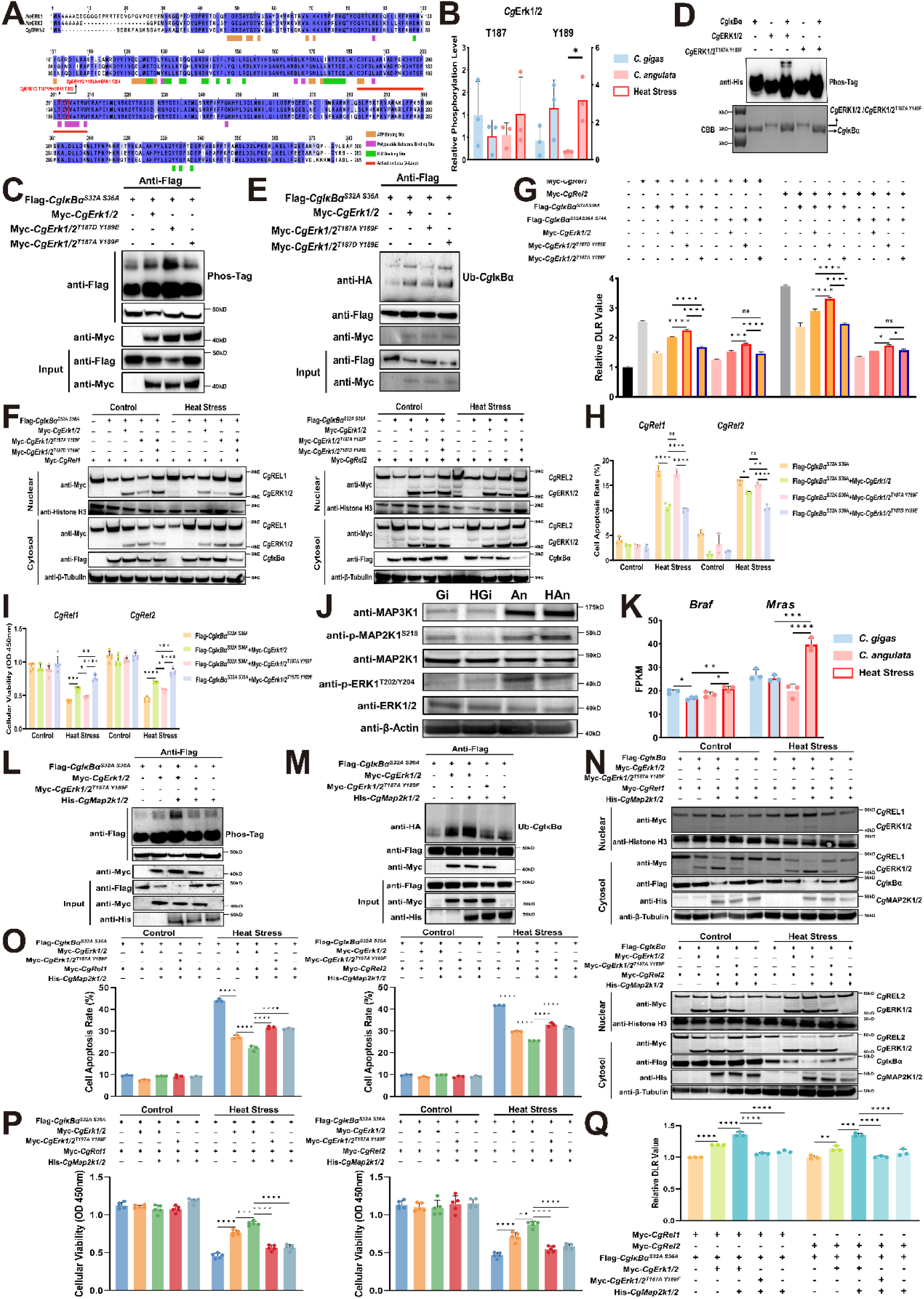
The MAPK/ERK signaling pathway regulates the phosphorylation level of Ser74 site at *Cg*IκBα. **A.** The sequence alignment of human ERK1, ERK2 and oyster ERK1/2. Different colors of sequence correspond to the degree of amino acid conservation (from light blue, low to blue, high). And the lower ribbon of sequence indicates different active sites or domain. Arrow marks the sequence alignment surrounding T187 and Y189 on *Cg*ERK1/2 protein. **B.** The phosphorylation levels of Thr187 and Tyr189 sites of *Cg*ERK1/2 in gill tissues of *C. gigas* and *C. angulata* during heat stress, which was obtained from our previous study (Wang et al., 2023a). **C.** *In vivo* phosphorylation assay of different *Cg*ERK1/2 mutants on *Cg*IκBα Ser74 site. HEK293T cells were transfected with Flag-*CgIκBα^S32AS36A^* and Myc-*CgErk1/2*/Myc-*CgErk1/2^T187AY189F^*/Myc-*CgErk1/2^T187DY189E^*. The cells were lysed and incubated with anti-Flag magnetic beads overnight, and IPed proteins were separated using SDS-PAGE gels containing Phos-tag (+Phos-tag). The phosphorylation level of Flag-*Cg*IκBα^S32AS36A^ was measured by western blotting using anti-Flag antibody. Protein samples were also separated in SDS-PAGE gels without Phos-tag (–Phos-tag). **D.** *In vitro* kinase activity assay of *Cg*ERK1/2/CgERK1/2^T187AY189F^ on *Cg*IκBα Ser74 site. The recombinant proteins (His-*Cg*ERK1/2/His-*Cg*ERK1/2^T187AY189F^ and His-*Cg*IκBα) were incubated in kinase buffer at 30 °C for 30 min and separated using SDS-PAGE gels containing Phos-tag, which were detected by western blotting with anti-His antibody. CBB, Coomassie Brilliant Blue staining. **E.** The protein ubiquitination experiments of *Cg*IκBα^S32AS36A^ co-transfected with *Cg*ERK1/2/*Cg*ERK1/2^T187AY189F^/*Cg*ERK1/2^T187DY189E^. The HEK293T cells were co-transfected with HA-Ub, Flag-*CgIκBα^S32AS36A^*and Myc-*CgErk1/2*/Myc-*CgErk1/2^T187AY189F^*/Myc-*CgErk1/2^T187DY189E^*, and incubated with 10 μM MG132 for 8 hours. Then, the cells were lysed and incubated with anti-Flag magnetic beads overnight, HA-tagged molecules co-IPed in this manner were resolved by SDS-PAGE and detected by immunoblotting with anti-HA antibody, which used to measure the ubiquitination level of *Cg*IκBα^S32AS36A^. **F.** The western blotting of nuclear and cytoplasmic proteins from HEK292T cells transfected with Flag-*CgIκBα^S32AS36A^*, Myc-*CgErk1/2*/Myc-*CgErk1/2^T187AY189F^*/Myc-*CgErk1/2^T187DY189E^*and Myc-*CgRel1*/Myc-*CgRel2* under control and heat treatment. **G.** The relative dual-luciferase reporter (DLR) values of HEK293T cells transfected with pNF-κB-Luc, pRL-TK, Myc-*CgRel1*/Myc-*CgRel2*, Flag-*CgIκBα^S32AS36A^*/Flag-*CgIκBα^S32AS36A^ ^S74A^*and Myc-*CgErk1/2*/Myc-*CgErk1/2^T187AY189F^*/Myc-*CgErk1/2^T187DY189E^*(n=3). The error bars represent the S.D. The cell apoptosis rate (**H.**;n=3) and the CCK-8 assay for the cell viability (**I.**; n=5) of HEK293T cells transfected with Flag-*CgIκBα^S32AS36A^*, Myc-*CgRel1*/Myc-*CgRel2* and Myc-*CgErk1/2*/Myc-*CgErk1/2^T187AY189F^*/Myc-*CgErk1/2^T187DY189E^*under control and heat treatment (n=3). The error bars represent the S.D. **J.** The western blotting of proteins extracted from gill tissues of *C. gigas* and *C. angulata* during heat stress with MAPK pathway’s antibodies. Gi and An represent the *C. gigas* and *C. angulata* under control condition. HGi and HAn represent the *C. gigas* and *C. angulata* under heat stress. **K.** The expressions of *CgBraf* and *CgMras* from transcriptomic data of *C. gigas* and *C. angulata* during heat stress (n=3) (Wang et al., 2024). **L.** *In vivo* phosphorylation assay of *Cg*IκBα^S32AS36A^ co-transfected with *Cg*ERK1/2/*Cg*ERK1/2^T187AY189F^ and *Cg*MAP2K1/2. HEK293T cells were transfected with Flag-*CgIκBα^S32AS36A^*, Myc-*CgErk1/2*/Myc-*CgErk1/2^T187AY189F^* and His-*CgMap2k1/2*. **M.** The protein ubiquitination experiments of *Cg*IκBα^S32AS36A^ co-transfected with *Cg*ERK1/2/*Cg*ERK1/2^T187AY189F^ and *Cg*MAP2K1/2. **N.** The western blotting of nuclear and cytoplasmic proteins from HEK292T cells transfected with Flag-*CgIκBα*, Myc-*CgErk1/2*/Myc-*CgErk1/2^T187AY189F^*, Myc-*CgRel1*/Myc-*CgRel2* and His-*CgMap2k1/2* under control and heat treatment. The cell apoptosis rate (**O.**; n=3) and the CCK-8 assay for the cell viability (**P.**; n=5) of HEK293T cells transfected with Flag-*CgIκBα^S32AS36A^*, Myc-*CgRel1*/Myc-*CgRel2*, Myc-*CgErk1/2*/Myc-*CgErk1/2^T187AY189F^* and His-*CgMap2k1/2* under control and heat treatment (n=3). The error bars represent the S.D. **Q.** The relative dual-luciferase reporter (DLR) values of HEK293T cells transfected with pNF-κB-Luc, pRL-TK, Myc-*CgRel1*/Myc-*CgRel2*, Flag-*CgIκBα^S32AS36A^*, Myc-*CgErk1/2*/Myc-*CgErk1/2^T187AY189F^*and His-*CgMap2k1/2* (n=3). The error bars represent the S.D. Significant differences among groups were marked with \**p*<0.05, \*\**p*<0.01, \*\*\**p*<0.001, and \*\*\*\**p*<0.0001. “ns” indicates non-significant differences.

Western blotting and expression experiments conducted on heat-stressed *C. gigas* and *C. angulata* demonstrated that *Cg*ERK1/2 phosphorylation levels were clearly higher in *C. angulata* than in *C. gigas* (Fig. 5J). Based on genome annotation and phylogenetic analysis, one Braf-and three Ras-like genes were found in oysters (Fig. S7A). Among them, the Ras-like genes were clustered into three subfamilies, including Mras, Rras2, and Ras-like. *CgRras2-* and *CgRas*-like genes showed no response to heat stress, except *CgMras* (Fig. S7B). The upstream regulators of ERK, including the phosphorylation level of MAP2K1/2 like (Fig. S6; Fig. 5J), relative expression of *Braf* (P<0.01; Fig. 5K) and *Mras* (P<0.001; Fig. 5K), and protein content of MAP3K1 (Fig. 5J) were significantly higher in HAn (*C. angulata* after heat stress) than in HGi (*C. gigas* after heat stress). The *in vivo* kinase experiment (Fig. 5L) and ubiquitination assay (Fig. 5M) proved that *Cg*MAP2K1/2 can increase the phosphorylation and ubiquitination levels of *Cg*IκBα^S32A^ ^S36A^ by phosphorylating *Cg*ERK1/2. Similarly, co-transfection of *Cg*MAP2K1/2 and *Cg*ERK1/2 caused an increased nuclear translocation of *Cg*REL in response to heat stress (Fig. 5N), significantly reducing cellular heat-induced apoptosis rate (P<0.0001; Fig. 5O) and promoting cell viability (P<0.0001; Fig. 5P). Furthermore, the luciferase reporter experiments confirmed that *Cg*MAP2K1/2 could enhance *Cg*ERK1/2 kinase activity through phosphorylation, promoting *Cg*IκBα^S32A^ ^S36A^ degradation and significantly increasing *Cg*REL-mediated transcriptional activity in the NF-κB signaling pathway (P<0.001; Fig. 5Q).

### Oyster’s MAPK/ERK/IκBα/REL regulatory axis exhibited environmental responsiveness

The 8-month-old F_1_ progeny of *C. gigas* and *C. angulata* were transplanted into their native and nonnative habitats, the northern (Qingdao, 35°44′ N) and southern (Xiamen, 24°33′ N) regions of China, and their corresponding protein phosphorylation and gene expression levels were evaluated after a 3-month reciprocal transplantation between local and non-local environments. The average seawater temperature in the southern habitat (19.64 ℃) was significantly higher than that in the northern habitat (11.24 ℃; P<0.0001, Fig. 6A), reflecting the distinct environmental temperature variations between the natural habitats of the two species. Western blotting of reciprocally-transplanted *C. gigas* and *C. angulata* demonstrated that the protein levels of *Cg*ERK1/2 and *Cg*MAP2K1/2 remained unaffected by the environment. However, their phosphorylation levels (*Cg*ERK1/2^T187^ ^Y189^ (corresponding to *Hm*ERK1^T202^ ^Y204^) and *Cg*MAP2K1/2^S238^ (corresponding to *Hm*MAP2K1^S218^)) showed a divergent pattern, with higher levels observed in the southern habitat than the northern habitat and in *C. angulata* than in *C. gigas* (Fig. 6B). Moreover, the protein content of CgMAP3K1 and gene expression levels of *CgBraf* and *CgMras* demonstrated a similar trend (Fig. 6C). Furthermore, the REL-mediated downstream genes, including *Abl*, *Elovl6*, *Acadm*, *Eif1b*, *Gst*, and *Map2k6* displayed the same expression pattern (Fig. 6D).

**Fig. 6.**
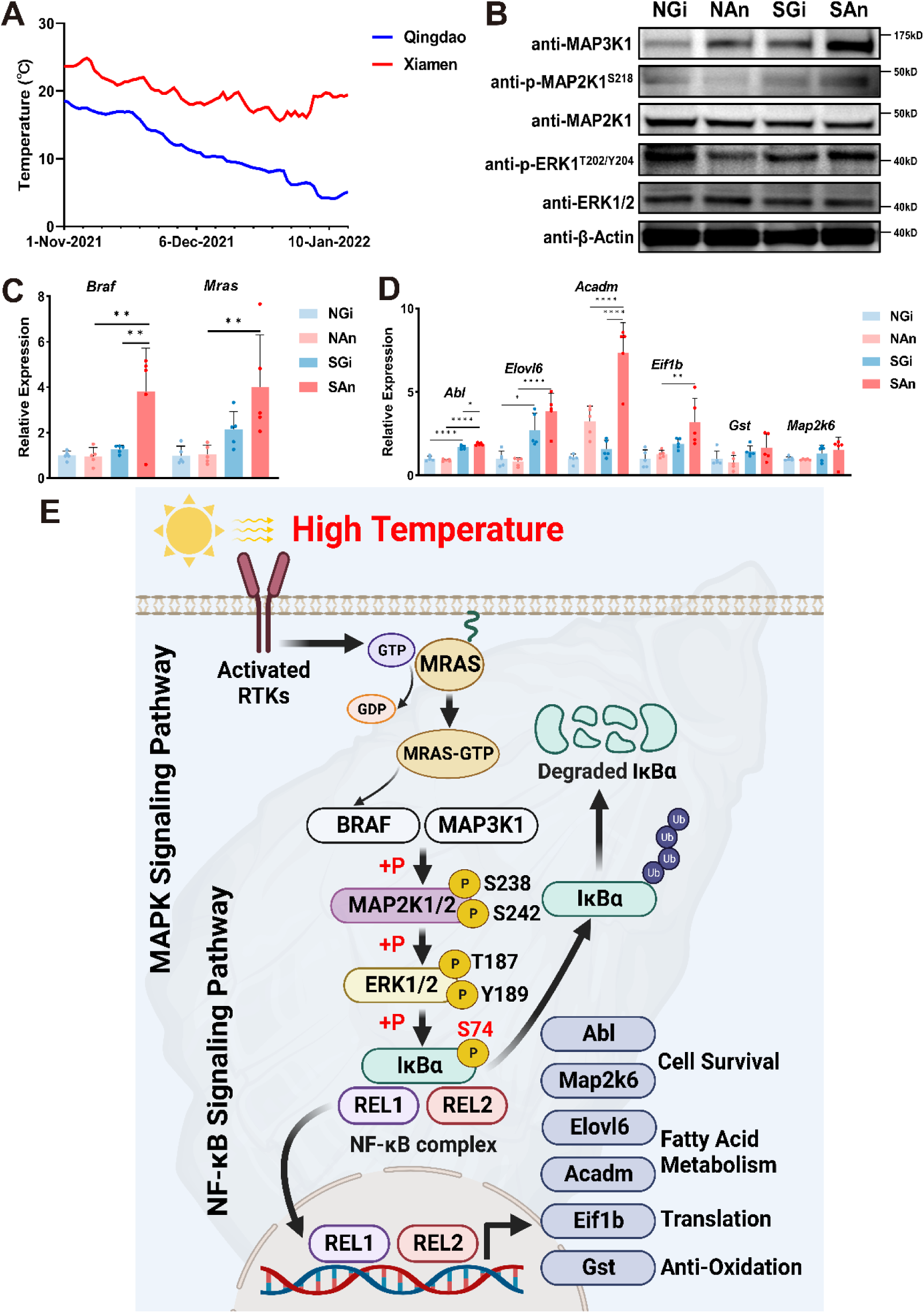
The MAPK/ERK/IκBα/REL axis demonstrates environmental responsiveness. **A.** Average Sea surface temperature of northern (Qingdao, 35°44′ N) and southern (Xiamen, 24°33′ N) sampling sites from November 2021 to January 2022. **B.** The western blotting of proteins extracted from gill tissues of *C. gigas* and *C. angulata* reared at northern (Qingdao, 35°44′ N) and southern (Xiamen, 24°33′ N) sampling sites. NGi and NAn represent the *C. gigas* and *C. angulata* were reared at northern site. SGi and SAn represent the *C. gigas* and *C. angulata* were reared at southern site. **C.** The relative expression of *CgBraf* and *CgMras* of *C. gigas* and *C. angulata* reared at northern (Qingdao, 35°44′ N) and southern (Xiamen, 24°33′ N) sampling sites. **D.** The relative expression of CgREL-mediated downstream genes (*CgBraf* and *CgMras)* of *C. gigas* and *C. angulata* reared at northern (Qingdao, 35°44′ N) and southern (Xiamen, 24°33′ N) sampling sites. **E.** Schematic representation of novel ERK/IκBα/REL axis of oyster in facing high temperature. Under thermal stress, the MAPK pathway is activated via RTKs-*Cg*MRAS complex. Then, the *Cg*MAP2K1/2 is phosphorylated by *Cg*BRAF or *Cg*MAP3K1 at Ser238 and Ser242 sites, which exerts kinase activity to phosphorylate *Cg*ERK1/2 at Thr187 and Tyr189 sits. Subsequently, *Cg*ERK1/2 phosphorylates *Cg*IκBα at the S74 site, facilitating its ubiquitin-proteasome degradation and releasing *Cg*REL1 and *Cg*REL2 for nuclear translocation, which as a transcription factor, activating genes involved in cell survival, fatty acid metabolism, protein translation, and antioxidant responses to assist oysters in combating heat stress.

## Discussion

The NF-κB signaling pathway, widely recognized as a conserved signaling pathway, has been extensively shown to be crucial in the biological immune processes of vertebrates and invertebrates (Ghosh & Hayden, 2008, Hayden & Ghosh, 2008, Kasthuri et al., 2013, Li et al., 2019, Sun & Ley, 2008, Xu et al., 2015). In this study, the phylogenetic analysis and sequence alignment identified three or four IκBα proteins in oysters, and most IκBα proteins in mollusks exhibit conserved Ser32 and Ser36 sites, consistent with the findings of previous studies (Montagnani et al., 2008, Sang et al., 2020, Xu et al., 2015). Notably, we identified a novel heat-induced phosphorylation site (Ser74) in the IκB protein of oysters. Of particular interest in this study is the *Cg*IκB like4 protein (*Cg*IκBα), which displayed the highest expression level among four IκBα proteins in *C. gigas* and possessed a specifically evolved Ser74 site unique to oysters. *Cg*IκBα demonstrated a response to high temperature and other abiotic stresses. Moreover, an opposite phosphorylation pattern at S74 (upregulated in *C. angulata* and downregulated in *C. gigas*) was observed between *C. gigas* (relatively low temperature adapted species) and *C. angulata* (relatively high temperature adapted species) during heat stress, whereas no phosphorylation modifications were detected at S32 and S36 (Wang et al., 2023a). Further functional experiments focusing on the phosphorylation of the *Cg*IκBα at the S74 site demonstrated that phosphorylation at this site can independently facilitate *Cg*IκBα degradation through the ubiquitin-proteasome pathway, which differs from the established mechanism in model organisms where phosphorylation at Ser32 and Ser36 sites by IKKs primarily controls the degradation of IκBα protein (Chen & Chen, 2013, Perkins, 2006). Additionally, the DSF results revealed that the phosphorylation at Ser74 of *Cg*IκBα significantly reduces its thermal stability, further proving that the decreased stabilization induced by phosphorylation can increase the ubiquitination level similar to p53 phosphorylation by aurora kinase A (Katayama, Sasai et al., 2004). Therefore, we propose that heat-induced novel phosphorylation at Ser74 of *Cg*IκBα can aid oysters in combating thermal stress by activating the NF-κB pathway through reduced thermal stabilization and increased ubiquitination and that its divergent heat response patterns in the two congenic oysters suggest its significant role in temperature adaptation. However, owing to the insufficient phosphoproteomics data in mollusks, we cannot determine whether phosphorylation at the Ser74 of *Cg*IκBα in oysters responds to other non-biological stressors.

Based on kinase prediction and subsequent functional validation, we confirmed that the MAPK/*Cg*ERK1/2 kinase phosphorylated Ser74 site of *Cg*IκBα, then activated the NF-κB/REL pathway to resist heat stress. ERK1 and ERK2 belong to the highly conserved MAPK family and phosphorylate many substrates, regulating a variety of evolutionarily conserved cellular processes in metazoans (Lavoie, Gagnon et al., 2020). Previous studies on the signal crosstalk between MAPK/ERK and NF-κB/IκBα pathways have found a significant association (positive (Chen, Yu et al., 2004, Funakoshi, Tago et al., 2001, Green, Macrae et al., 2011, Lin, Tang et al., 2012, Liu, You et al., 2006) or negative (Maeng, Min et al., 2006, Yeh, Yeh et al., 2004)) between the phosphorylation levels of ERK and IκBα, which are usually mediated through IKKs.However, our study first reported a direct kinase-substrate interaction between MAPK/ERK and NF-κB/IκBα proteins in oysters exposed to high-temperature, which bypasses the known IκBα upstream kinase IKKs and does not rely on the Ser32 and Ser36 sites. Further measurements of phosphorylation and expression levels in key kinases and regulatory factors of the classical MAPK pathway showed that *C. angulata* demonstrated a stronger activation pattern in classical MAPK pathway than *C. gigas* during heat stress and under increased wild environmental temperature, as evidenced by higher phosphorylation levels of *Cg*ERK1/2 at T187 and Y189 and *Cg*MAP2K1/2 at S238 and S242, the protein content of MAP3K1, and the expression levels of *CgBraf* and *CgMras*. MAPK/ERK pathway reportedly participates in the high-temperature stress response in marine invertebrates (Anestis, Lazou et al., 2007, Evans & Somero, 2010, He, Xiong et al., 2023, Liu, Li et al., 2023, Xu, Wang et al., 2023) and other taxa, inducing the expression of heat shock proteins (Keller, Escara-Wilke et al., 2008, Yang, Zhao et al., 2021), mitigating oxidative stress and mitochondrial damage (Wang, Wu et al., 2023c, Wu, Ibtisham et al., 2019), and ultimately promoting cell survival and inhibiting apoptosis (Siddiqui, Khan et al., 2023). The co-transfection of *Cg*MAP2K1/2 and *Cg*ERK1/2 demonstrated that activated *Cg*ERK1/2 can further upregulate the phosphorylation at the S74 site of *Cg*IκBα protein and subsequent degradation, increasing the transcriptional activity of *Cg*REL. According to the existing classical MAPK pathway, the activation of ERK1/ERK2 is achieved through the phosphorylation of Thr-202/185 and Tyr-204/187 sites by MAP2K1/MAP2K2 (Kinoshita, Yoshida et al., 2008), the Ser-218/222 and Ser-222/226 sites of MAP2K1/MAP2K2 are phosphorylated by RAF or MAP3K1 (Lavoie, Sahmi et al., 2018), and the RAF is activated by Ras-GTP through membrane recruitment and other kinases (such as PKA, PAK, and SRC) (Yao, Zhang et al., 1995). The RAS family can be divided into four types—KRAS, NRAS, HRAS, and MRAS. The MRAS exhibited high-temperature responsiveness in this study and is reportedly crucial in enhancing MAPK signal transduction by counteracting inhibitory phosphorylation events on the RAF protein family through SHOC2-MRAS-PP1C complex (Kwon, Hajian et al., 2022). Additionally, there is evidence suggesting that RAS can be activated by the receptor tyrosine kinase (RTK) to regulate cell proliferation and survival through the downstream MAPK cascade during heat stress, similar to those induced by growth factor stimulation (Lin, Hu et al., 1997). Our previous study revealed significant differences in the protein content and phosphorylation modifications of RTKs, such as FGFR3, during the thermal stress between *C. gigas* and *C. angulata* (Wang et al., 2023a). Our results indicate that the presence of a high-temperature-RTK-MRAS-BRAF-MAPK-NF-κB signaling cascade in oysters, with the core components being the activation of MAPK/ERK and the phosphorylation of its substrate IκBα (Ser74), followed by ubiquitin-mediated proteasomal degradation, which exhibits inter-species differentiation and responses to elevated environmental temperatures, and in essential in shaping the divergent temperature adaptation.

The function experiments had proven that NF-κB/*Cg*IκBα interact with *Cg*REL1 and *Cg*REL2 in the cytoplasm, suppressing its nuclear translocation and transcriptional activity, which are consistent with previous reports on the other three *Cg*IκBα proteins (Montagnani et al., 2008, Xu et al., 2015, Zhang, He et al., 2011). Subsequent investigations had revealed that *Cg*REL possessed a high-temperature nuclear translocation capability, which is crucial in cellular and oysters’ defenses against apoptosis induced by heat stress. This is consistent with the findings by Liu et al. that the levels of RelA (p65) and phosphorylation of IκBα in chickens subjected to high temperature significantly increased, supporting the involvement of NF-κB/Rel pathway in the response to heat stress (Liu, Ding et al., 2022). Using the divergent phosphorylation levels of *Cg*IκBα S74 under heat stress between *C. gigas* and *C. angulata*, in conjunction with DAP-Seq, transcriptome following RNAi experiment, and transcriptomic and ATAC-Seq data in *C. gigas* and *C. angulata* during heat stress obtained from previous study, along with experimental validations of regulatory relationship and binding affinity, we successfully identified six differentially *Cg*REL-activated genes between two species during high temperature. *Abl*, a pivotal tyrosine protein kinase, is crucial in cell survival processes such as cytoskeleton remodeling, DNA damage response, and cell apoptosis (Gu, Lavau et al., 2012, Wie, Adwan et al., 2014). *Map2k6* activates the p38MAPK signaling cascade, regulating enzyme activity (MK2 and MSK1) and transcription factor expression (p53 and ATF2), which mediate cellular survival responses under stress conditions (Chan, Hsu et al., 2005, Gutiérrez-Uzquiza, Arechederra et al., 2012, Shi, Jin et al., 2011). Fatty acids, as essential sources and structural materials, have been extensively studied in the high-temperature stress condition (Iba, 2002). *Acadm*, encoding an acyl-CoA dehydrogenase, participates in the initial step of mitochondrial fatty acid beta-oxidation, providing energy for thermal injury repair (He, Pei et al., 2011). Moreover, *Cg*REL regulates *Elovl6*, a key enzyme in fatty acid metabolism that catalyzes the initial rate-limiting reaction in the elongation cycle of long-chain fatty acids, consequently impacting the synthesis of saturated fatty acids, such as C16:0 (Ohno, Suto et al., 2010). An increased ratio of saturated/unsaturated fatty acid is reportedly involved in the resistance to heat stress by regulating cell membrane fluidity, otherwise termed homeoviscous adaptation (Mendoza, 2014). Our previous studies have also found that *C. angulata* demonstrates a higher degree of fatty acid saturation (Wang et al., 2023b, Wang et al., 2021) and a correspondingly higher heat-responsive capacity of saturated fatty acid synthesis genes (Wang, Jiang et al., 2024). Ribosomal proteins have emerged as significant markers for physiological status evaluation under stress conditions in marine organisms (Jiao, Dai et al., 2021, Vieira, Braga et al., 2018). *Eif1b* participates in the assembly of the 43S complex during the mRNA translation of protein synthesis (Sun, Gong et al., 2015), remodeling the protein landscape to enhance stress survival during heat stress (Neef & Thiele, 2009), which is consistent with our previous findings (Wang et al., 2023a). Furthermore, *Gst*, a well-known antioxidant gene, shows increased expression and enzyme activity in marine organisms under high-temperature stress, aiding in combating oxidative stress induced by elevated temperatures (Guerriero, Bassem et al., 2018, Missionário, Travesso et al., 2023, Teixeira, Diniz et al., 2013). *C. angulata* has been proven to possess a stronger anti-oxidative capacity than *C. gigas* under heat stress (Wang et al., 2023a). The differential expression patterns of the above genes in *C. gigas* and *C. angulata* under heat stress and in the North and South environments further support the activation of the MAPK/ERK-NF-κB/IκBα cascade in response to high temperature, and *C. angulata* possesses stronger activation of this cascade, which activates the expression of cellular survival, fatty acid metabolism, protein translation, and antioxidant-related genes, participating oysters’ temperature adaptation.

To conclude, the results of this study demonstrated the existence of a unique high temperature-induced crosstalk mechanism between the MAPK and NF-κB pathways in oysters. A schematic of the signal cascade of high temperature-RTK-MRAS-BRAF-MAPK-NF-κB pathways in oysters during heat stress is provided in Figure 6E. This mechanism involved direct phosphorylation of novel Ser74 site at major *Cg*IκBα by MAPK/*Cg*ERK1/2, bypassing the Ser32 and Ser36 sites phosphorylated by IKK proteins, independently causing its ubiquitin-proteasome degradation and decreased thermal stability. The NF-κB pathway was regulated by the classical MAPK pathway, which exhibited stronger activated patterns in response to higher environmental temperature in *C. angulata* than in *C. gigas*, two allopatric congeneric oyster species with differential habitat temperatures. Finally, this signaling cascade activated cell survival, fatty acid metabolism, protein translation, and antioxidant gene expression by promoting *Cg*REL nuclear translocation to resist heat stress. These findings highlight the presence of complex and unique phosphorylation-mediated regulatory networks in marine invertebrates, which differ from the established mechanisms in vertebrates and contribute to expanding our understanding of the evolution and function of the crosstalk mechanisms between existing classical pathways.

## Methods

### Animal Material

Wild adult oyster of *C. gigas* and *C. angulata* were collected from their natural habitat in Qingdao (35°44′ N) and Xiamen (24°33′ N), respectively, and used as broodstock (Wang et al., 2023b). The one-generation common garden experiment was conducted to alleviate the environmental effects (Sanford & Kelly, 2011). Artificial breeding program including broodstock conditioning, fertilization, and larval cultures, all of which were conducted in hatchery with 22-26 °C and 31 ± 1 ‰ seawater. Briefly, the eggs of 30 mature females were mixed and divided into 30 beakers for each species, each fertilized with sperm from one of the 30 males, to maximize parental contribution. Juvenile F_1_ progeny (8 months old) of each species were separated into two groups: one group was deployed at the northern site (35°44′N, Qingdao, Shandong province, China) and the other group was deployed at the southern site (24°33′N, Xiamen, Fujian Province, China). Three wo months later, 10-month-old oysters were sampled from each site or collected to laboratory for subsequent studies. The sea water at two sites was recorded using HOBO Conductivity U24 Data Loggers with a time interval of 2 h during November 2021-January 2022.

### Phylogenetic Analysis

Each member of human IκB family was retrieved from the National Center for Biotechnology Information (NCBI) database. Using TBtools Blast with an E-value threshold of 10^−5^, a BLASTP analysis was performed to identify all possible NF-κB proteins in the oyster and other species’ amino acid sequence databases (Chen, Chen et al., 2020). Then, the selected proteins were subjected to ANK domain identification using SMART (Simple Modular Architecture Research Tool) (http://smart.emblheidelberg.de) software. And the non-IkB family proteins were manually removed based on results of domain identification above. The maximum likelihood phylogenetic tree was reconstructed using PhyloSuite 1.2.2 software (Zhang, Gao et al., 2020), incorporating the inferred amino acid sequences of all putative NF-κB family genes in invertebrates, as well as the ERK gene in oysters, and representative genes from vertebrates such as humans. Sequences alignments were conducted using PRALINE software (http://www.ibi.vu.nl/programs/pralinewww/), and poorly aligned sequences and gaps were removed using Gblocks 9.1b (http://www.phylogeny.fr/one_task.cgi?task_type=gblocks). The optimal models were tested by ModelFinder (Kalyaanamoorthy, Minh et al., 2017), and the LG+G4 model (NF-κB) and LG+I+G4 model (ERK) were selected for multi-species tree reconstruction. IQ-TREE integrated into PhyloSuite was used for maximum likelihood tree construction with 1000 bootstrapping replicates (Minh, Schmidt et al., 2020). See Table S1 for details.

### Oysters IκB Gene Expression Analysis

The raw sequencing data of the transcriptomes of oysters from different tissues, developmental stages, and under different stress conditions were obtained from Short Read Archive (SRA) (http://www.ncbi.nlm.nih.gov/sra) database (SRP014559; SRP019967) (Zhang et al., 2012, Zhang, Li et al., 2015). The data were aligned to the updated genome of *Crassostrea gigas* (GenBank accession no. GCA_011032805.1) (Qi, Li et al., 2021) and calculated using Salmon software (https://combine-lab.github.io/salmon/) (Patro, Duggal et al., 2017). Heatmap was generated using the OmicShare tools (https://www.omicshare.com/tools).

### Molecular Evolutionary Analyses and Ancestral Reconstruction

The PAML 4.10.6 software package (Yang, 2007) and its user interface PAMLX (Xu & Yang, 2013) were used to analyze the levels of selective pressure acting upon the Mollusca IκBα proteins. Firstly, the PAML branch model was employed to investigate whether the specific branch containing the IκBα proteins with the conserved S74 site in oyster is under selection in mollusks. The one-ratio model assumes a common dN/dS ratio for all branches, while the two-ratio model assumes that the branch of interest has a different ratio of nonsynonymous to synonymous substitutions (dN/dS; ω1) compared to the background ratio (dN/dS; ω0). Then, the random sites models (M1a, M2a, M3, M7, M8, M8a) implemented in the CODEML program were used to estimate the evolutionary rates (dN/dS) at site 179 (*C. gigas* IκBα like4 S74) within mollusk IκBα proteins. Significant differences in model fits were determined by likelihood ratio tests.

### Co-IP

Co-IP assays were conducted using the Flag-tag Protein IP Assay Kit with Magnetic Beads (Beyotime Biotechnology, China). The full-length CDSs of *CgIκBα*, *CgRel1*, *CgRel2* and *CgErk* were amplified and inserted into pCMV-N-Flag and pCMV-N-Myc plasmids for fusion with the tag (Table S3). The Lipofectamine 3000 (Invitrogen, USA) was used to transfect these plasmids into HEK293T cells (Procell Life Science & Technology, China), which was cultured in high-glucose DMEM Medium (Biological Industries, Israel) and 10% fetal bovine serum (Biological Industries, Israel). The Co-IP reaction was performed according to the manufacturer’s instructions. Briefly, the cells lysates were incubated with anti-Flag magnetic beads and Mouse IgG magnetic beads overnight. Following washing step, the reaction products were loaded onto 4∼20% SDS-PAGE gels (GenScript Biotech, China), and then the signals were obtained by western blotting.

### Subcellular Localization

The full-length CDSs of *CgIκBα*, *CgRel1*, *CgRel2* and *CgErk* were amplified and inserted into pCMV-N-EGFP, pCMV-N-mCherry and pCMV-N-CFP plasmids for fusion with the reporter gene, respectively (Beyotime Biotechnology, China). The Lipofectamine 3000 (Invitrogen, USA) was used to transfect these plasmids into HeLa cells (Procell Life Science & Technology, China), which was cultured in RPMI Medium 1640 (Biological Industries, Israel) and 10% fetal bovine serum (Biological Industries, Israel). After incubation at 42 °C for 2h, the fluorescence was imaged using a confocal microscope LSM710 (Carl Zeiss, Germany).

### BiFC Assay

The full-length CDSs of *CgIκBα*, *CgRel1*, *CgRel2* and *CgErk* were amplified and inserted into pBiFC-VN173 and pBiFC-VC155 plasmids (MiaoLing Plasmid Platform, China; Table S3). The cell culture, plasmid transfection and imaged for fluorescence as mentioned in the section “Subcellular Localization”.

### Luciferase Reporter Assay

The promoter region of *CgRel* downstream genes was amplified and inserted into pGL3-Basic plasmid (MiaoLing Plasmid Platform, China; Table S3). The pRL-TK Renilla luciferase plasmid, pNF-κB-luc (Beyotime Biotechnology, China) or recombinant pGL3 plasmid and different plasmids combinations were transfected into HEK293T cells as above.

Luciferase activity was measured using the Dual-Luciferase Reporter Assay System (Promega, USA) and measured using Varioskan Flash multimode reader (Thermo Fisher Scientific, USA). All experiments were conducted three technical replicates, and the firefly luciferase activity was normalized to the Renilla luciferase activity of each sample.

### Nuclear and Cytoplasmic Isolation

The cells were treated with 2h of 42 °C, and then lysed using NE-PER Nuclear and Cytoplasmic Extraction Reagents (Thermo Fisher Scientific, USA). Then, the nuclear and cytoplasmic proteins were loaded onto 4∼20% SDS-PAGE gels and detected using western blotting.

### Immunohistochemistry

Rabbit polyclonal antibodies against *Cg*REL1 and *Cg*REL2 were generated by Wuhan Bqbio Biological Technology, China. Immunohistochemical staining was performed according to following protocol. The gill tissues of *C. gigas* and *C. angulata* under control and heat stress treatments were fixed and embedded by paraffin (Servicebio, China). Then, the sections were de-waxed and rehydrated. Antigen was retrieved using Tris-EDTA antigen retrieval solution (PH9.0) at 37 °C for 15 min (Servicebio, China). The solution of 3% H_2_O_2_ was used to block the activity of endogenous peroxidase. Antibodies to *Cg*REL1 and *Cg*REL2 were added and incubated overnight at 4 °C. HRP-conjugated goat anti-rabbit secondary antibody was then added and incubated for 50 min at room temperature, following by color development with a DAB Chromogen Kit (Servicebio, China). Nuclei were shown with hematoxylin counterstain (Servicebio, China).

### EMSA

The recombinant plasmids pCMV-N-Myc encoding *CgRel1* and *CgRel2* were transfected into HEK293T cells as above. Cells transfected with pCMV-N-Myc plasmid were utilized as a control, and the cellular nuclear proteins were extracted for the ESMA experiment. The biotin-labeled EMSA probe specific to NF-κB and its corresponding unlabeled cold probe were acquired from Beyotime Biotechnology, China. The DNA probe labeled with 5′ biotin was synthesized by Tsingke Biotechnology, and biotinylated and unlabeled probe sequences were as follows: F, 5′-GGGCAATTAC-3′. For super-shift assay 1ug of anti-Myc antibody (ABclonal, China) was added. Binding reactions, and detection of probe shifts were performed using the LightShift® Chemiluminescent EMSA Kit (Thermo Fisher Scientific, USA) according to the manufacturer’s instructions.

### Cell Apoptosis Assay

Cell apoptosis after 2h of 42 °C was analyzed using Annexin V-fluorescein isothiocyanate (FITC) and propidium iodide (PI) apoptosis detection kit (Solarbio, China) according to the manufacturer’s instructions. And the apoptotic cells were detected using the FACSAria II flow cytometry (Becton Dickinson, USA). Data was analyzed using FlowJo (version 10.8.1).

### Cell Counting Kit-8 (CCK-8) Assay

Cell death after heat stress (42 °C, 2h) was measured using the Enhanced Cell Counting Kit-8 (Beyotime Biotechnology, China) at 36 h after transfection, according to the manufacturer’s instructions. The Varioskan Flash multimode reader (Thermo Fisher Scientific, USA) was used to measure the absorbance at 450 nm.

### RNAi Experiment

Small interfering RNA (siRNA) was synthesized by GenePharma (Shanghai, China) and used for the RNAi experiment (sequences of the siRNA were shown in Table S2). The RNAi experiment was based on our previous study (Wang et al., 2023b). Briefly, oysters (*C. gigas*) were cleaned and reared in 500 L tank for one week. Then, individuals were anesthetized (500 g MgCl2+5 L seawater +5 L freshwater) and injected with 100 μl of 10 μg/100 μl siRNA (siRNA group, n=15), 10 μg/100 μl NC strands (NC group, n=15) and 100 μl seawater (water group, n=15). Based on the results of the pilot experiment, we selected the most effective siRNA and interference time. In the formal experiment, gill tissue was sampled from 15 individuals in each group at 24 h post-injection for subsequent RNA extraction and qRT-PCR. Meanwhile, a continuous heat stress experiment was performed at 42°C (sublethal temperature), and the mortality of oysters was recorded hourly. Survival curves were analyzed and plotted based on the Kaplan-Meier method using GraphPad Prism version 8.0.2 for Windows.

### RNA-seq

RNA extraction from gill tissues of *C. gigas* in RNAi experiment was performed using TRIzol reagent (Tsingke Biotechnology, China). Every five samples were pooled in equal quantities to form a single biological replicate. The NEBNext Ultra™ RNA Library Prep Kit for Illumina (NEB, USA) was used for generating the RNA-Seq library for each sample, which was then subjected to sequencing on the NovaSeq 6000 platform. To ensure the acquisition of high-quality and clean reads, Trimmomatic software (v0.36) (Bolger, Lohse et al., 2014) was implemented to eliminate adapter sequences, reads containing 10% of unknown nucleotides (N), and reads with more than 50% low-quality bases (Q-value ≤ 20). Subsequently, the clean reads were aligned to the oyster genome (GenBank accession no. GCA_011032805.1) (Qi et al., 2021) using HISAT2 v2.1.0 (Kim, Langmead et al., 2015). The HTSeq v0.6.0 tool was used to count the reads for each gene in each sample to quantify gene expression (Anders, Pyl et al., 2015). The DESeq2 R package (v1.6.3) (Love, Huber et al., 2014) was utilized to identify statistically significant changes in gene expression (q-value ≤ 0.05 and |log2(fold change)| ≥ 1). The sequencing data in this study has been deposited into the Sequence Read Archive (SRA) BioProject, under the accession number PRJNA1059049.

### DAP-Seq

The full-length CDSs of *CgRel1* and *CgRel2* were ligated with pFN19K HaloTag® T7 SP6 Flexi® Vector using ClonExpress II One Step Cloning Kit (Vazyme Biotech, China). Gene Denovo Biotechnology Co. (Guangzhou, China) submitted recombinant vectors for the in vitro expression of candidate proteins and conducted DNA affinity purification sequencing (DAP-seq) (Bartlett, O’Malley et al., 2017). The genomic DNA was extracted from the gill tissue of *C. gigas*. The DNA was then sonicated to generate fragments of approximately 200 bp in length, followed by the construction of a DNA library using magnetic bead-based size selection. Recombinant proteins were subsequently immobilized on magnetic beads and mixed with the DNA library. Eluents were then used for PCR amplification and bead purification. The resulting products were sequenced using the Illumina HiSeq™ 4000. Raw reads were processed to obtain clean reads by removing reads containing adapters and low-quality reads, which were then aligned to the oyster genome (GenBank accession no. GCA_011032805.1) (Qi et al., 2021) using the Bowtie2 software (v2.2.5) (Langmead & Salzberg, 2012). DeepTools (v3.2.0) was employed to determine read counts within the intervals spanning from transcription start sites (TSS) to transcription end sites (TES), as well as the upstream and downstream 2-kb intervals (Ramírez, Ryan et al., 2016). The DAP-seq data was subsequently analyzed using the MACS2 software (v2.1.2) (Zhang, Liu et al., 2008) to identify regions of enriched reads. The dynamic Poisson distribution was used to calculate *p*-value based on the read counts from uniquely mapped reads within specific regions. Regions with a q-value of 0.05 were considered as peaks. The ChIPseeker R package was employed to annotate the peak-related genes (Yu, Wang et al., 2015). The distribution of peaks across genomic regions, including intergenic, introns, downstream, upstream, and exons was analyzed. The sequencing data in this study has been deposited into the Sequence Read Archive (SRA) BioProject, under the accession number PRJNA1058704.

### Differential Scanning Fluorimetry (DSF)

The recombinant purified proteins, His-*Cg*IκBα and His-*Cg*IκBα^S74A^ were utilized for performing Differential Scanning Fluorimetry (DSF) experiment to assess the influence of Ser74 site phosphorylation on protein thermal stability. DSF was performed using 10 μM protein and 10X SYPRO Orange (Invitrogen, USA). The samples were subjected to a heating ramp from 25°C to 95°C at a rate of 1°C/s using the ABI 7500 Fast Real-Time PCR System (Applied Biosystems, USA). Data was analyzed by Protein Thermal Shift Software v1.4 to obtain melting temperature (Tm).

### *In vivo* Ubiquitination Assay

The *in-vivo* ubiquitination assay was utilized to assess the degree of ubiquitination of the *Cg*IκBα protein at various phosphorylation levels. HEK293T cells were co-transfected with Flag-*CgIκBα* plasmid or its corresponding site-mutated variants, as well as the HA-Ub plasmid (MiaoLing Plasmid Platform, China). Following an 8-hour treatment with MG132 (10 μM; MCE, USA), the ubiquitinated IκBα protein was purified using the anti-Flag magnetic beads as mentioned in the section “Co-IP”. The purified product was subsequently separated by 4-20% SDS-PAGE and subjected to western blotting.

### Yeast Two-Hybrid Assay

The full-length CDSs of *CgIκBα* and *CgErk1/2* were amplified and inserted into pGBK-T7 and pGAD-T7 (MiaoLing Plasmid Platform, China), respectively (Table S3). Pairwise interactions were tested using GAL4 Yeast Two-Hybrid Media Kit (Coolaber, China). Briefly, each vector (bait and prey) was transformed in the Y2HGold yeast strain, and plated first on -Leu -Trp plates to allow selective growth of transformants. After 2-3 days, growth transformants were inoculated into (-Leu, -Trp) medium and allowed to grow overnight at 30°C with continuous shaking at 200 rpm. Subsequently, 10 μl of cell suspension, diluted in ddH_2_O to achieve an optical density (OD) of 0.5 and 1.0, was plated on selective plates (-Leu, -Trp), (-Leu, -Trp, -His3) and (-Leu, -Trp, -His3, -Ade2) and incubated for 2-3 days to assess the presence of interactions.

### Prediction of kinase

The iGPS1.0 software was utilized for predicting potential kinases targeting the *Cg*IκBα S74 site. This prediction was based on the Short Linear Motif (SLM) theory, which focuses on the surrounding short linear motifs associated with the phosphorylation site (p-site), providing a high degree of specificity (Song, Ye et al., 2012). For the analysis, *Homo sapiens* was chosen as the organism, with a threshold set to "medium" and the "interaction" parameter configured as "Exp./String".

### *In vivo* Phosphorylation Assay

Recombinant Flag-*CgIκBα* plasmid and its site-mutated variants, along with additional upstream plasmids such as Myc-*CgErk1/2* plasmid, were co-transfected into HEK293T cells as above. After 36 h, the cells were lysed using a lysis buffer supplemented with protease and phosphatase inhibitors (Beyotime Biotechnology, China). The IκBα protein was subsequently purified using the anti-Flag magnetic beads as mentioned in the section “Co-IP”. The phosphorylation levels of the IκBα protein were then measured using the Phos-tag SDS-PAGE method. Phos-Tag™ was purchased from FUJIFILM Wako Pure Chemical Corporation, Japan and used according to the manufacturer’s protocol. Briefly, gels used for Phos-tag SDS-PAGE consisted of a separating gel [8% (w/v) acrylamide, 375 mM Tris-HCl, pH 8.8, 50 μM Phos-tag acrylamide, 100 μM MnCl_2_, 0.1% SDS solution, 0.1% (v/v) N,N,N’,N’-tetramethylethylenediamine (TEMED), and 0.05% (w/v) ammonium persulfate (APS)], and a stacking gel [4.5% (w/v) acrylamide, 125 mM Tris-HCl, pH 6.8, 0.1% SDS solution, 0.1% (v/v) N,N,N’,N’-tetramethylethylenediamine (TEMED), and 0.05% (w/v) ammonium persulfate (APS)]. Electrophoresis was conducted at a constant current of ≤30 mA/gel with the running buffer [25 mM Tris-base, 192 mM glycine, and 0.1% (w/v) SDS]. For western blotting analysis, gel was washed using wash buffer [25 mM Tris, 192 mM glycine, 0.1% (v/v) SDS, 10 mM EDTA] for 10 min three times to remove metal ions, followed by one wash without EDTA for 10 min. Then, the samples were electroblotted to the polyvinylidene difluoride (PVDF) membrane (Millipore, USA) for subsequent western blotting.

### *In Vitro* Kinase Activity Assay

The full-length CDSs of *CgIκBα* and *CgErk1/2* and their site-mutated variants, were amplified and inserted into pET-32a plasmid to express the 6 X His-tags at both N- and C-ends (Table S3). The purified proteins were used to test whether recombinant CgERK1/2 could phosphorylate CgIκBα, as well as to evaluate the influence of phosphorylation at the T187 and Y189 sites of CgERK12 on its ability to phosphorylate CgIκBα. The wild type and S74A mutant of CgIκBα was used as substrate. The kinase activity assay was performed in 20 μl kinase buffer containing 25 mM Tris-HCl, pH 7.5, 5 mM beta-glycerophosphate, 2 mM dithiothreitol (DTT), 0.1 mM Na_3_VO_4_, 10 mM MgCl_2_, 20mM ATP (CST, USA), 1 μg substrate protein and 1 μg kinase protein for 30 min at 30 °C. And the reactions were stopped with SDS loading buffer and electrophoresed on Phos-tag SDS-PAGE as mentioned in the section “*In vivo* Phosphorylation Assay”.

### Raw Data Reanalysis

The transcriptomic, ATAC-Seq, and protein phosphorylation omics data for *C. gigas* and *C. angulata* under 12 h of heat stress were downloaded from our previous study. The ATAC-Seq signals in target region were calculated using DeepTools (v3.2.0) (Ramírez et al., 2016) and visualized using the Integrated Genomic Viewer (version 2.16.0) (Thorvaldsdóttir, Robinson et al., 2013). The heatmap were visualized using using the OmicShare tools (https://www.omicshare.com/tools).

### qRT-PCR Experiment

Gill tissues from oyster were collected for total RNA extraction using the TRIzol reagent (Tsingke Biotechnology, China). First-strand cDNA was generated using HiScript III RT SuperMix for qPCR (Vazyme Biotech, China). The primers were designed using Primer 5 software and synthesized by Tsingke Biotechnology (detailed sequences of primers were shown in Table S4). The Ef-1α gene was used as an internal control. The qPCR was conducted using the Taq Pro Universal SYBR qPCR Master Mix (Vazyme Biotech, China) in the ABI 7500 Fast Real-Time PCR System (Applied Biosystems, USA). The 2^−△△CT^ method was used to calculate the relative expression (Livak & Schmittgen, 2001). The detailed comparison between the two species has been previously described (Wang et al., 2023b).

### Western Blotting

The samples from oysters and cells were extracted using Cell lysis buffer for Western and IP (Beyotime Biotechnology, China) and M-PER™ Mammalian Protein Extraction Reagent containing protease inhibitor (Thermo Fisher Scientific, USA) supplemented with protease and phosphatase inhibitors (Beyotime Biotechnology, China), respectively. The supernatant protein was collected by centrifugation, and then subjected to denaturation at 100 °C for 10 min after the addition of 4X protein loading buffer (GenScript Biotech, China). Then, the proteins were transferred onto 0.45 nm pore polyvinylidene fluoride (PVDF) membrane (Millipore, USA) using an eBlot™ L1 wet transfer (GenScript Biotech, China). Membranes were blocked and incubated with primary antibodies and secondary antibodies using eZwest Lite Automated Western Device (GenScript Biotech, China). Membranes were then incubated with Omni-ECL™Femto Light Chemiluminescence Kit (Epizyme, China) and captured using the Molecular Imager® Gel Doc™ XR System (Bio-Rad, USA). The antibodies used were as follows: Flag-tag (ZENBIO, 390002), Myc-tag (ZENBIO, 390003), beta Tubulin (ZENBIO, 200608), Histone H3 (ZENBIO, R24572), beta Actin (ZENBIO, 200068-8F10), HA-tag (ZENBIO, 201113), His-tag (ZENBIO, 230001), ERK1/ERK2 (ABclonal, A16686), Phospho-ERK1- T202/Y204 + ERK2-T185/Y187 (ABclonal, AP0472), MAP2K1/MAP2K2 (ABclonal, A24394), Phospho-MAP2K1-S217/MAP2K2-S221 (ABclonal, AP0209), MAP3K1 (ABclonal, A21490), HRP-labeled Goat Anti-Mouse/Rabbit IgG(H+L) (Epizyme, LF101) and HRP-labeled Goat Anti-Mouse/Rabbit IgG(H+L) (Epizyme, LF102).

### Statistical Analysis

All statistical analyses were performed using GraphPad Prism version 8.0.2 for Windows. After confirming the normality of the distributions using the Shapiro–Wilk test and homogeneity of variance using Bartlett’s test, data were analyzed with the two-tailed unpaired Student’s t-test, one-way analysis of variance (ANOVA) and two-way ANOVA followed by Tukey’s multiple comparisons test. Data are shown as the means ± SD, and the number of replicates (n) are denoted in the corresponding figure legends. Significant differences between groups were marked with “*” for P< 0.05, “**” for P<0.01, “***” for P<0.001 and “****” for P<0.001. The schematic presentation was created using BioRender software (https://biorender.com).

## Acknowledgements

The authors would like to thank the supercomputer cluster of the High-Performance Computing Center (HPCC) at the Institute of Oceanology, Chinese Academy of Sciences for support in bioinformatics analysis.

## Author contributions

L.L. and G.Z. conceived the study. C.W., Z. J., M.D., T.Z. and J.C carried out the field and laboratory work, collected the oyster samples, participated in the data analysis, and drafted the manuscript. R.C. and W.W. contributed to cultural management. C.W., L.L. and G.Z. revised the manuscript. All authors approved the manuscript for publication.

## Data availability

The raw sequencing data of transcriptome in RNAi experiment and DAP-Seq in this study have been deposited the Sequence Read Archive (SRA) BioProject under the accession number: PRJNA1059049 and PRJNA1058704.

